# VerteBrain reveals novel neural and non-neural protein assemblies conserved across vertebrate evolution

**DOI:** 10.1101/2025.05.26.656196

**Authors:** Vy Dang, Brittney Voigt, David Yang, Gabriel Hoogerbrugge, Muyoung Lee, Rachael M. Cox, Ophelia Papoulas, Claire D. McWhite, Raksha Pradeep, Janelle C. Leggere, Benjamin A. Neely, Ryan S. Gray, Edward M. Marcotte

**Author notes:** Correspondence to: E.M.M.

## Abstract

Protein-protein interactions underlie core brain functions, including neurotransmitter release, receptor activation, and intracellular signaling essential for learning, memory, and cognition. Here, we systematically map conserved brain protein interactions across five vertebrate species–rabbit, chicken, dolphin, pig, and mouse–using co-fractionation and immunoprecipitation mass spectrometry. From 2,197 biochemical fractions, we identify over 81,000 high-confidence interactions among 6,108 conserved proteins. This interaction map (VerteBrain) reveals both regulatory and structural complexes, including extensive synaptonemal protein associations likely involved in inter-neuronal coordination. Conservation across species underscores essential roles in neuronal and glial function, as well as in additional tissues for more widely expressed complexes. The VerteBrain dataset uncovers candidate disease mechanisms, including roles for ARHGEF1 in short stature syndromes, synaptic vesicle trafficking complexes in epilepsy, and RELCH in congenital deafness. VerteBrain provides a publicly accessible framework for investigating brain protein interactions and their relevance to human neurological disorders.

## INTRODUCTION

Protein-protein interactions (PPIs) form the basis of nearly every cellular process, enabling precise coordination of proteins within complex biological systems. In the vertebrate brain, PPIs are particularly important, especially for neuronal communication at synapses, where PPIs are responsible for the clustering of receptors that mediate signaling, ultimately regulating synaptic transmission, plasticity, and memory formation. While proteomic analyses have yielded valuable insights into brain function, a comprehensive understanding of how brain proteins are organized into multi-protein complexes remains elusive. Disruptions in these assemblies underlie diverse neurodevelopmental disorders, underscoring the need to systematically define brain protein interactions. Moreover, as estimates suggest 80-95% of human genes are expressed at some stage in the human brain^1^, brain PPIs should also broadly inform studies in other tissues. Mapping these assemblies should thus help guide our understanding of the molecular logic of brain and other tissue function, as well as provide a framework for interpreting the impact of disease-linked mutations, and ultimately facilitate targeted treatments.

Over the last decade, researchers have generated comprehensive transcriptomic and connectomic brain atlases, providing invaluable insights into brain biology at the cellular level and the etiology of diseases^2–5^. In contrast, proteomic atlases, particularly those capturing protein-protein interactions, remain sparse. This is primarily due to the proteome’s inherent complexity: proteins are dynamic, often extensively post-translationally modified, and exist within specialized spatial and temporal contexts, all of which introduce technical challenges. Global neuroproteomic studies also face additional challenges due to the heterogeneity and complexity of brain tissues and the corresponding difficulties of capturing endogenous protein interactions on a large-scale.

Despite such challenges, recent advances in proteomics have made it possible to systematically survey protein interactions in complex systems. Various *in vitro* methods from yeast two-hybrid (Y2H) assays^6^ to recently developed orthogonal mass spectrometry-based techniques have been applied to define the interactomes of various biological systems. To date large-scale (high-throughput) physical protein interaction maps have been reported for systems ranging from yeast^7^, plants^8^, human cell lines^9,10^, to specific cell-types such as red blood cells^11^. For the brain, the largest interaction datasets have been measured from postmortem mouse^12,13^ and human brain^14^ tissues, as well as from specific compartments such as mouse synaptosomes^15^ (recently reviewed in Dang & Voigt *et al*.^16^). While these studies offer key insights, comprehensive brain interactomes remain limited, especially given the difficulties in accessing and preserving healthy human brain tissue, suggesting there is an important opportunity for leveraging the relative ease of obtaining healthy tissue from evolutionary diverse animals in order to better define conserved brain interactions. Such proteomic data could then be integrated with transcriptomic and connectomic atlases to enhance our understanding of brain biology and uncover new pathways relevant to neurological disease.

To address the need for more comprehensive brain protein interaction networks, we applied Co-Fractionation Mass Spectrometry (CF-MS), an ultra-high-throughput technique that enables proteome-wide mapping of endogenous protein interactions without the need for antibodies or genetic tagging, unlike related techniques such as affinity purification mass spectrometry (AP-MS). CF-MS can thus be applied broadly across species, making it uniquely suited for comparative interactomics, especially in non-model organisms, as it requires only orthogonal biochemical separations, high resolution mass spectrometry, and supervised machine learning. We applied CF-MS to native soluble protein extracts from the postmortem brains of four vertebrate species–chicken, dolphin, pig, and rabbit–integrating these new datasets with previously published mouse brain CF-MS experiments^12,13^. We focused exclusively on evolutionarily conserved proteins and interactions, as conservation significantly increases confidence that these interactions are critical to brain function. This study reports the resulting cross-species interactome of the vertebrate brain: VerteBrain.

To validate our observed interactions, we conducted extensive immunoprecipitation mass spectrometry (IP-MS) analyses and experimental validation in vertebrate model organisms. Notably, we observed protein complexes active both in the brain as well as in non-neural tissues, underscoring the broader biological relevance of the detected assemblies. While one of the ultimate goals for neuroscience research is to develop therapeutics for debilitating neurological diseases, access even to healthy brain tissue remains inherently limited. VerteBrain demonstrates that comparative analysis across evolutionary diverse species can yield robust insights into conserved mechanisms of brain function and their disruption in disease.

## RESULTS AND DISCUSSIONS

### A comprehensive dataset of brain protein abundance and purification profiles from five vertebrates

We generated a rich proteomics dataset from 5 different vertebrate species spanning over 319 million years of evolution: *Mus musculus* (mouse), *Sus scrofa* (pig), *Oryctolagus cuniculus* (rabbit), *Gallus gallus* (chicken), and *Tursiops erebennus* (Tamanend’s bottlenose dolphin) (**Figure 1A**). We performed 20 in-house biochemical fractionations directly on native protein lysates from whole brain tissues from chicken and dolphin, as well as dissected brain regions from pig and rabbit brains (frontal cortex, cerebellum, hippocampus and temporal lobe). Together with the incorporation of mouse brain CF-MS data from previously published studies^12,13^, the data represent mass spectrometry of 2,197 native biochemical fractions from post-mortem brains of chicken, dolphin, pig, rabbit, and mouse, (**Figure 1B**).

**Figure 1.**
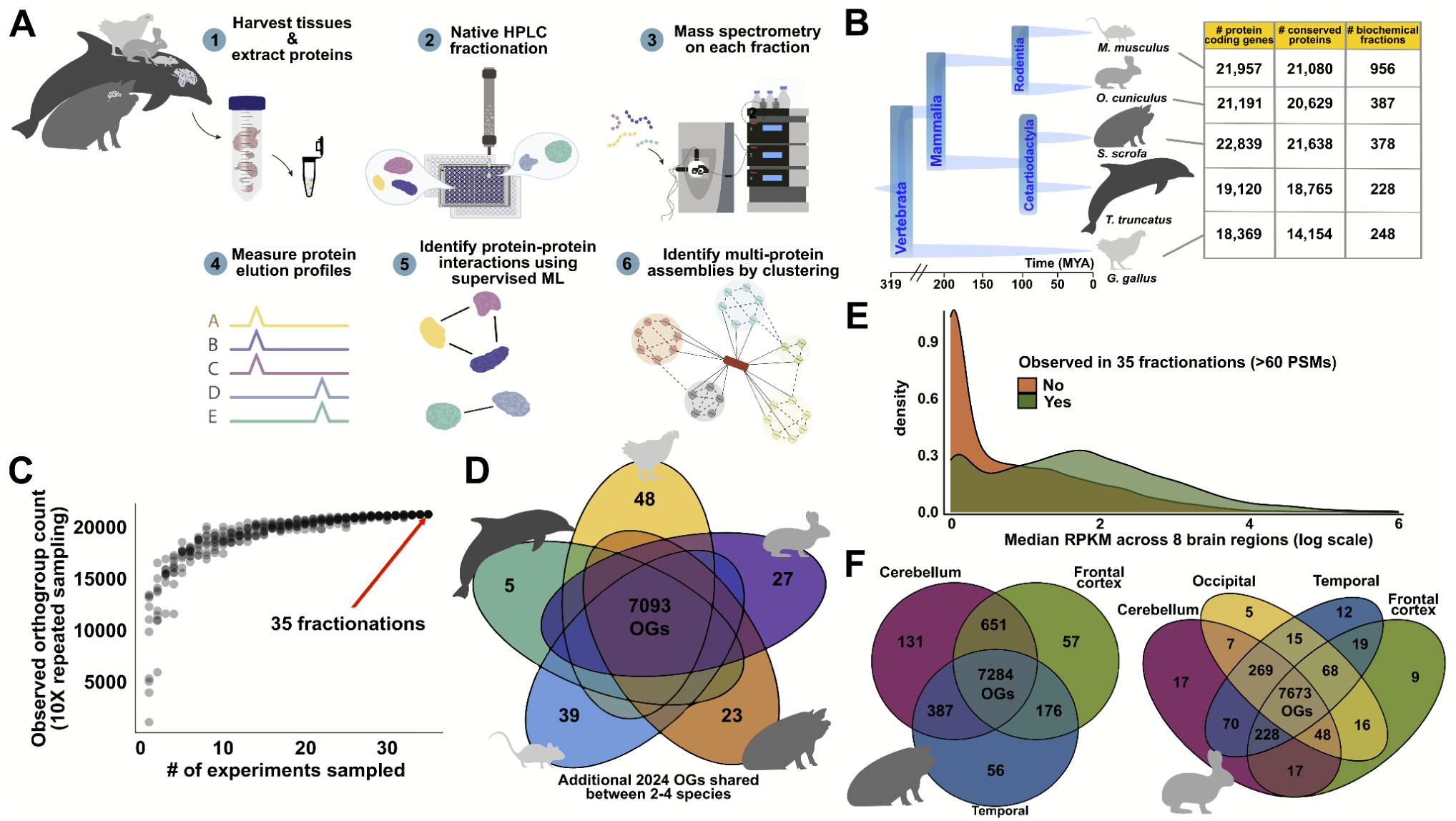
Overview of methods, samples, data collection, and proteome coverage. (A) Schematic of Co-Fractionation Mass Spectrometry experimental and computational workflows. (B) Phylogenetic relationship of the species used in the study, their respective proteome sizes, and the number of mass spectrometry experiments performed for each species. (C) The total number of orthogroups detectable by mass spectrometry is saturated by 35 CF-MS experiments used in the study. Each dot represents the number of OGs identified (y-axis) in a subsample of n experiments (x-axis), with sampling repeated ten times per n to evaluate saturation. (D) A core set of 7,093 conserved OGs is detected across all 5 species. (E) Proteins detected in biochemical fractionation exhibit a shift to higher median RPKM values (log scale), suggesting that the proteins analyzed are not only well-conserved but also functionally active in diverse brain regions. (F) A core set of 7673 OGs is detected across all tested distinct brain regions of rabbit and pig.

We separated each native, non-denatured protein extract by some biophysical property, either size exclusion chromatography (SEC), ion exchange chromatography (IEX), or weak anion exchange chromatography (WAX). Each chromatographic fraction was analyzed by high-resolution, high-sensitivity liquid chromatography (LC-MS). We carried out replicate runs of our separations to ensure reproducibility. Each species contributed at least 200 mass spectrometry experiments, ensuring a reasonably balanced representation across the species (**Figure 1B**). Further description of experimental details of species, tissues and fractionation procedures can be found either in the **Methods** section or in **Table S1**. Across all datasets, we identified 14,653,734 interpretable peptide mass spectra (sum of all peptide spectral matches (PSMs) across all protein groups and fractions). This deep proteome profiling dataset thus captures brain proteins across vertebrates, serving as a valuable resource for investigating neurobiological questions and understanding the evolution of synaptic proteins in the vertebrate nervous system.

In order to consider these data in a comparative proteomics framework, we mapped proteins to their corresponding vertebrate orthologous groups. We adopted a strategy similar to McWhite *et al*.^8^ and Cox *et al.*^17^, matching mass spectral observations to orthogroups rather than individual proteins. An orthogroup (OG) is defined as a set of orthologous genes—genes in different organisms that originate from a single gene in their last common ancestor, which often encode proteins performing the same function across species. We considered orthogroups at the vertebrate level as calculated by the eggNOG^18^ (evolutionary genealogy of genes: non-supervised orthologous groups) database of orthology relationships, which provided the framework for comparing proteomics data across vertebrates and ultimately to human gene annotations. This orthogroup-based proteomics approach allows peptides shared among multiple proteins within an OG, but not unique to a single protein, to contribute to quantification. The spectra were subjected to stringent database searching and filtering (false discovery rate <1% at both peptide– and protein-level) using three different MS-search algorithms^19^.

Overall, our approach resulted in high coverage of proteins and notably, the 35 CF-MS experiments were sufficient to saturate the detection of OGs (**Figure 1C**). To focus downstream analyses on well-observed proteins, we required a minimum abundance for each orthogroup of 60 PSMs from the union of all fractions. In terms of human proteins, the resulting filtered dataset covers 11,025 unique proteins belonging to 9,259 Vertebrata OGs, sampled from extracts of brain tissues of five vertebrate species.

Several metrics indicate that the dataset is of high quality. First, OGs already known to be associated with neurodegenerative disorders and disease, or essential for neurogenesis, modulation of neurotransmitter release at the pre-synaptic terminal, or maintenance of dendritic architecture^20^ are among the highest in abundance. These include microtubule-associated protein 2 (MAP2)^21^, dihydropyrimidinase-related protein 2 (DPYSL2)^22^, spectrin alpha chain, non-erythrocytic (SPTAN1), and the synapsin protein family (SYN1–SYN3).

Second, we investigated the extent to which high abundance OGs from the experiments are detected across all species. Indeed, over 75% (7,093 of 9,259) high-abundance OGs are observed across all 5 species (**Figure 1D**), underscoring the evolutionary support within this dataset. Quantification of the full set of 9,259 high abundance OGs can be found in **Table S2**.

Third, we asked to what extent peptide level data in VerteBrain correspond to the transcript level data identified in the RNA-seq dataset of eight distinct human brain regions from the Adult Genotype Tissue Expression Portex (GTEx) v8.0. We found that vertebrate proteins robustly observed in CF-MS experiments (at least 60 PSMs across 35 fractionation experiments) were more likely to correspond to higher mRNA expression levels across eight distinct human brain regions (**Figure 1E**). This finding highlights the nuanced and imperfect correlation between protein-level detection and transcriptional activity^23,24^, even on a cell-by-cell basis^25^, while also reinforcing the reliability of our dataset. Finally, we examined the diversity of proteins recovered from different brain regions within the same species, and observed a core set of proteins observed broadly across different brain regions (**Figure 1F**). Our CF-MS strategy thus yielded a high-quality, deeply sampled proteomic dataset from vertebrate brains, with strong representation of neurobiologically relevant proteins, extensive detection across species and brain regions, and broad but only partial correspondence with human brain transcriptomic data.

### Defining a well-conserved interactome among vertebrate brain proteins

In many cases, well-characterized and stable molecular machines exhibited clustered elution patterns that could easily be detected by eye. Examples include the 19S regulatory subunits of the proteasome, the Commander holo complex, and the actin-related protein 2/3 complex (**Figure 2A**). However, a robust computational framework is required to systematically identify these patterns in a high-throughput manner and effectively control for false positives. To address this, we implemented a supervised machine learning strategy, similar to approaches used in previously published studies of large-scale interactomes^8,11,17^. This method leverages the fact that known protein complexes are observed within the separations and can serve as internal standards, and assigns a probabilistic CF-MS score between 0 and 1 to each observed PPI, where a score of 1 represents the highest level of confidence that a pair of proteins is interacting and 0 indicates no evidence of interaction.

**Figure 2.**
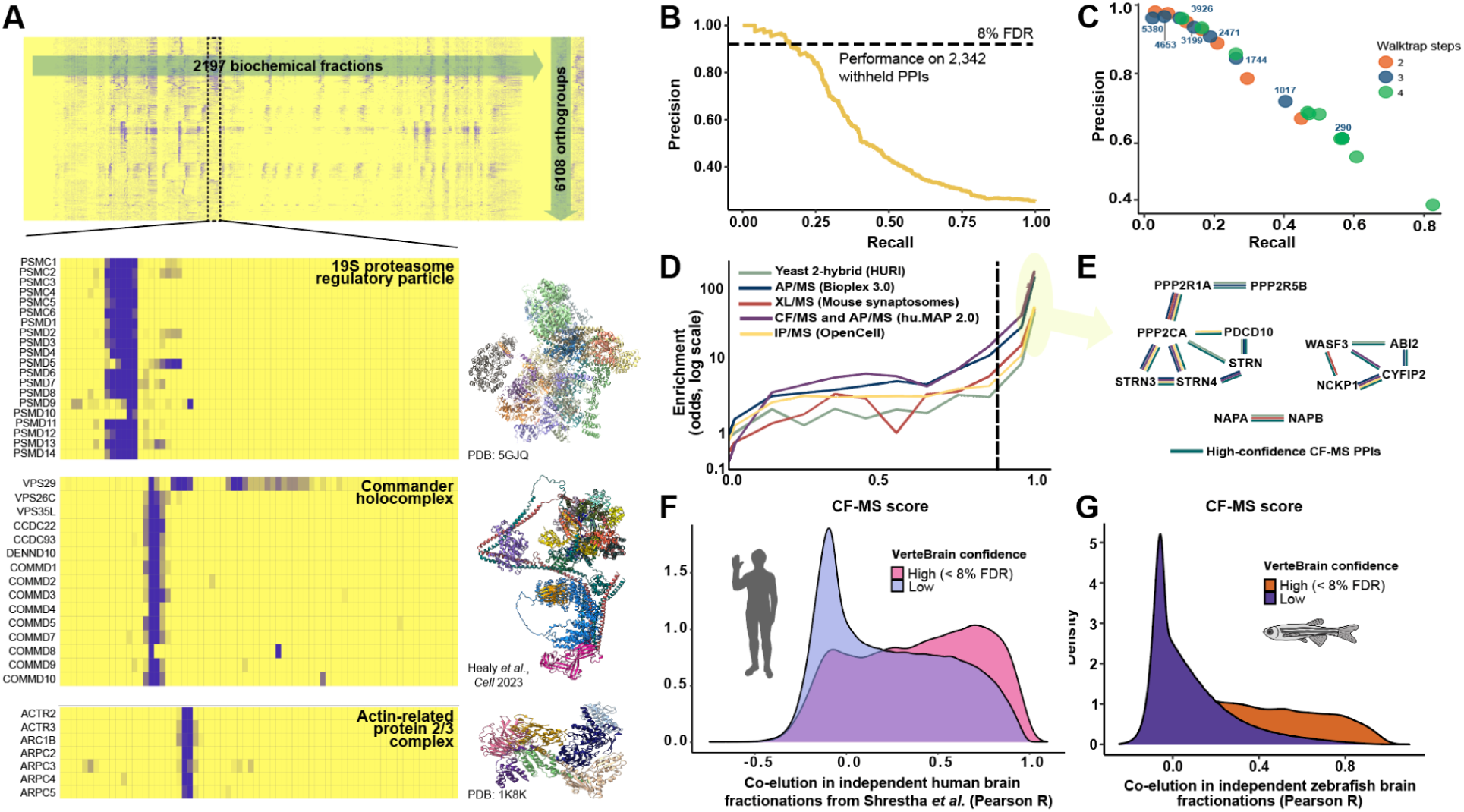
Identifying protein interactions from the CF/MS datasets and validating with independent experiments. (A) Heatmap of the full dataset of abundance measurements for 9259 well sampled OGs across 2,197 fractionations for 5 species. (B) Precision-recall performance of 2,342 withheld PPIs. (C) Precision-recall curves for the reconstruction of known protein complexes defined by a walktrap algorithm, where pairwise PPI scores from each number of walks (colored as in legend) are used as input. Points are labeled with the total number of clusters (complexes) created at each point in the hierarchy. (D) VerteBrain shows strong agreement with orthogonal techniques that also measure PPIs. Dotted line indicates the 8% FDR score cutoff (CFMS score ≥ 0.88). (E) Agreement of CF-MS PPIs pairwise interactions (dark green) with Y2H (light green), AP/MS (dark blue), XL/MS (red), huMAP2.0 (purple), and IP/MS (yellow) for three example protein complexes. (F) PPIs with high confidence CF-MS scores (≥ 0.88) in VerteBrain are highly correlated with the CF-MS dataset of human brain published by Shrestha *et al.*^14^ (G) PPIs with high confidence CF-MS scores (≥ 0.88) in VerteBrain are highly correlated with the independent CF-MS dataset measured from zebrafish brains.

In order to train the classifier and score the pairwise interactions, we first computed pairwise similarities between proteins’ elution profiles, considering a variety of measures and normalizations (see **Methods**). The resulting numerical measures served as features for a supervised classification algorithm, selected using automated machine learning^26^ to best recover a training set of known protein complexes as defined by the CORUM protein complex database^27^, a compendium of complexes manually curated from the literature. We observed the best performance by using an ExtraTreeClassifier with 5-fold cross-validation and a set of the 70 best-performing features. We evaluated the model’s performance using a test set of 2,342 known pairwise PPIs that were fully withheld from model training (**Figure 2B**). For interactions with CF-MS scores exceeding 0.88, the model achieved 92% precision and 18% recall (see Zenodo repository for data), demonstrating excellent accuracy for high-confidence interactions. Out of the 12,961,622 pairs of proteins evaluated by the model (see **Methods**), 81,225 PPIs had scores above the 0.88 threshold, representing high-confidence protein interactions. In total, the resulting brain protein interactome included 6,108 OGs, each with at least one interaction above the 8% FDR threshold based on the ExtraTreeClassifier model. Pairwise interactions are provided in **Table S3**.

### Defining and validating multi-protein assemblies in VerteBrain

To define coherent protein complexes from the high-confidence pairwise interactions, we employed the unsupervised community detection algorithm walktrap^28^, which has previously been shown to perform well for protein complex identification^11,17^ (see **Methods**). This algorithm reweights pairwise interactions and organizes them into a dendrogram, which can then be cut at specific depths to define a nested hierarchy of complexes. Depending on the granularity of the cutoff, the algorithm identifies 145 (less granular) and 5380 (more granular) protein “neighborhoods” (**Figure 2C**), which at finer granularity correspond strongly to discrete protein complexes. This hierarchical organization thus enables exploration of both broad and fine-grained protein interactions, facilitating insights into the functional organization of the brain interactome. The set of protein assemblies defined in this way is reported in **Table S4**.

We wanted to know how well our large-scale measurement of PPIs aligns with external datasets. To this end, we benchmarked CF-MS scores against independent interactome studies that used orthogonal techniques to measure PPIs (see **Methods**). Overall, PPIs with high CF-MS scores were more likely to be detected by Y2H assays (in HuRI^7^), AP-MS experiments (in BioPlex 3.0^10^), chemical cross-linking (XL-MS) in mouse synaptosomes^15^, integration of AP-MS and CF-MS (in hu.MAP 2.0^29^), and by live-cell fluorescent microscopy coupled with immunoprecipitation mass spectrometry (as recorded in the OpenCell^30^ database), when compared to randomly paired proteins (**Figure 2D**). Notably, PPIs surpassing the 8% FDR threshold displayed a very strong correlation with other methods, strongly supporting the accuracy of our reported PPIs.

As an independent test to see if high CF-MS scores indicate more frequent direct contacts between the interacting proteins, we integrated our CF-MS data with 3D structure prediction using AlphaFold-Multimer^31^. This approach not only enabled us to validate previously established PPIs but also to provide additional support for novel PPIs. By integrating structural modeling with CF-MS data, we aimed to elucidate the structural basis of their interactions where possible. We performed an *in silico* PPI screen using AlphaFold-Multimer on protein pairs identified in the VerteBrain dataset, grouping them into bins based on their CF-MS scores (in 0.1 increments). We then examined the relationship between CF-MS score bins and AlphaFold-Multimer’s interface predicted template modeling (ipTM) score, which estimates the accuracy of the predicted relative positions of interacting subunits. Overall, we observed that protein pairs in score bins exceeding the FDR threshold exhibited higher AlphaFold confidence scores (**Figure S1**). However, while high ipTM scores have been shown to be reliable indicators of accurate protein complex predictions across multiple studies^32–34^, it should be noted that low ipTM scores (i.e., < 0.25) do not necessarily mean that two proteins do not interact. For example, subunits of the same protein complex may not directly contact each other, but they will still co-elute in CF-MS experiments and may have low ipTM scores in AlphaFold-Multimer predictions. This is because they are part of the same larger structure and are therefore physically associated, even if they do not share a binding interface.

It is also important to note that no single technique captures all PPIs perfectly, as each method has their inherent strengths and weaknesses. For instance, although all techniques are capable of capturing the majority of well established pairwise interactions among members of the evolutionary conserved STRIPAK complex (<u>str</u>iatin-interacting <u>p</u>hosphatase <u>a</u>nd <u>k</u>inase)^35^, only Y2H, AP-MS, and CF-MS detected the interaction between PDCD10 (<u>P</u>rogramme<u>d</u> <u>c</u>ell <u>d</u>eath protein <u>10</u>) and Striatin (STRN) (**Figure 2E**). This discrepancy may be explained by the differential expression pattern of the striatin protein family: STRN is highly enriched in the CNS, STRN4 is predominantly expressed in the brain and lung, while STRN3 is ubiquitously expressed across all tissues^36,37^. Additionally, VerteBrain demonstrates a remarkable capacity to detect neural-specific protein interactions across a broad spectrum of structural and functional categories. Examples range from large, multisubunit assemblies like the WAVE (<u>WA</u>SP family <u>Ve</u>rprolin homolog—also known as SCAR for <u>S</u>uppressor of <u>cA</u>MP <u>R</u>eceptor) regulatory complex, to simpler heterodimeric interactions, such as the NSF attachment proteins alpha (NAPA) and beta (NAPB) (**Figure 2E**). Some interactions, including those identified through our CF-MS observations, can only be captured by specialized techniques, such as Y2H or targeted studies of specific cellular compartments, like mouse synaptosomes. This highlights the complementary nature of various PPI mapping approaches and reinforces the unique strengths of VerteBrain for uncovering context-specific and neural-enriched protein networks.

### Cross-species validation confirms reliability and biological relevance of VerteBrain PPIs

As an additional test of the PPIs identified, we compared VerteBrain to two species-specific PPI maps. First, we observed the same overall trend in CF-MS data from a human brain study published by Shrestha *et al.*^14^ (**Figure 2F**). As a further test of conservation across vertebrates, we conducted two additional CF-MS experiments on adult zebrafish brain lysates using SEC and IEX. To ensure the reliability and quality of our zebrafish CF-MS data, we examined internal controls by analyzing the elution profiles of well-characterized protein complexes. Specifically, we confirmed that all subunits of the CCT (<u>c</u>haperonin-<u>c</u>ontaining <u>T</u>CP-1) chaperone and the COP9 (<u>Co</u>nstitutive <u>p</u>hotomorphogenesis <u>9</u>) signalosome complex co-eluted as intact assemblies across both SEC and IEX experiments, validating the integrity and resolution of our chromatographic fractionation (**Figure S2**). We observed that high confidence VerteBrain PPIs tended to exhibit highly correlated elution profiles in the zebrafish brain data, supporting the generality of our interaction across vertebrates (**Figure 2G**). These analyses strengthen the confidence in our CF-MS workflow and provide additional support for the biological relevance of the detected interactions.

### VerteBrain recapitulates the known organization of many neuronal proteins

In order to assess the biological relevance of our dataset, we evaluated VerteBrain’s recovery of well-characterized multiprotein assemblies critical for neuronal function. As illustrated for representative examples in **Figure 3A**, we examined a diverse array of essential neuronal processes, spanning multiple regions of the nervous system, including the heterotetramic Clathrin Adaptor Complex-2 (AP-2)^38^, which plays a pivotal role in clathrin-dependent endocytosis, a process essential for retrieving synaptic vesicles at presynaptic terminals following neurotransmitter release. In neurons, AP-2 is also critical for neurodevelopment by regulating cell polarity during differentiation^39^ and promoting neuronal survival^40^, axon growth cone outgrowth and guidance^41^, as well as dendrite formation and extension^40,42^. VerteBrain confidently identified high scoring pairwise interactions between AP-2 subunits (e.g., AP2A1/AP2A2, AP2B1, and AP2M1), and also recapitulated other broadly functioning complexes, such as the prefoldin complex and the tRNA ligase complex. Beyond these large, well-studied assemblies, VerteBrain also detected more specialized complexes, such as the neurofilament (NEFL, NEFM, and NEFH) involved in maintaining neuronal architecture^43^, and the Post-Synaptic Density Adapters, such as the SHANK family (<u>SH</u>3 and Multiple <u>Ank</u>yrin Repeat Domains), which play critical roles in synaptic signaling and organization. Additionally, VerteBrain confidently recovers interactions within the SNARE (<u>s</u>oluble <u>N</u>SF <u>a</u>ttachment protein <u>re</u>ceptors) complex, a key regulator of vesicle transport and neurotransmitter release^44^. These examples illustrate the breadth of the dataset in uncovering known ubiquitous and neuron-specific assemblies.

**Figure 3.**
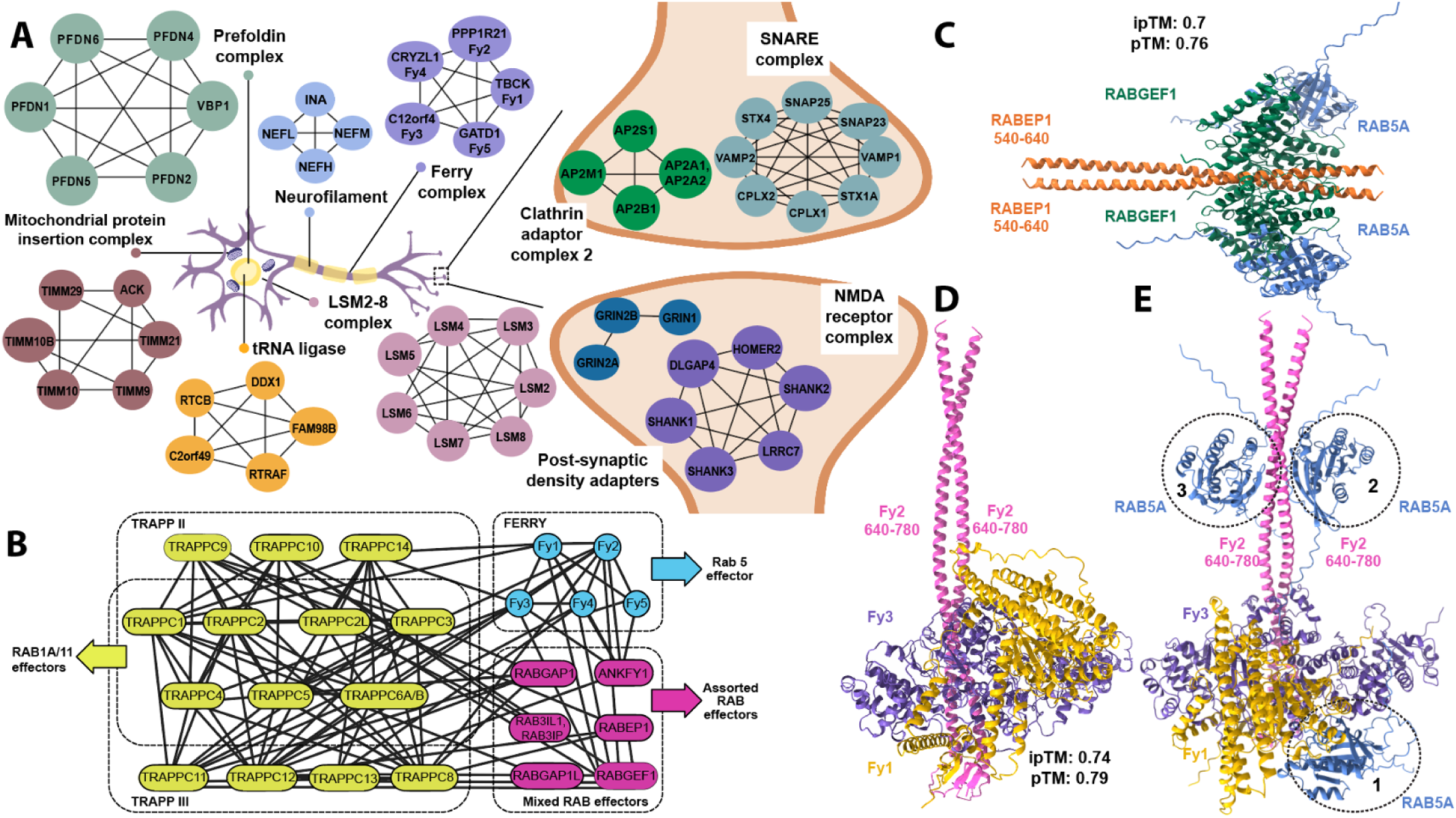
VerteBrain recapitulates many known neuronal protein complexes and interactions. (A) VerteBrain analysis identified numerous well-characterized neuronal protein complexes spanning a range of sizes, including the NMDA receptor complex, prefoldin complex, clathrin adaptor complexes, and additional assemblies essential for neuronal function. (B) Detected interactions encompassed subunits of the TRAPPII and TRAPPIII complexes, as well as crosstalk among established Rab effectors. (C) AlphaFold3 multimer modeling predicted the most favorable stoichiometry for the RABEP1 (residues 540–640; orange)–RABGEF1 (dark green)–RAB5A (blue) complex to be 2:2:2. (D) AlphaFold3 multimer model is in well agreement with schematic representation the of FERRY complex by Quentin *et al.*^53^, capturing Fy1 (yellow) and Fy3 (purple) binding to a homodimer of Fy2 (residues 640–780; pink), consistent with hydrogen/deuterium exchange mass spectrometry data. (E) Predicted partial structure of the FERRY complex highlights three potential Rab5A (blue) binding sites, with a suggested order of preferential binding from site 1 to 3.

We also more systematically measured VerteBrain’s reconstruction of known systems by quantifying the extent to which VerteBrain clusters proteins known to function together. Specifically, we calculated the area under the receiver operating characteristic curve (AUROC) for proteins known to belong to the same Gene Ontology (GO) in the same GO protein-containing complex (PCC) or cellular anatomical entity (CAE) (**Figure S3**). As expected, random protein sets yielded AUROC scores centered around 0.5, indicating no predictive power. In contrast, VerteBrain demonstrated significantly higher AUROC scores for proteins with shared GO term annotations, confirming the biological coherence and robustness of the dataset. Similar to previously reported large-scale interactome studies on brain tissues^12–14^, our dataset showed significant enrichment for protein networks associated with the cytoskeleton, extracellular matrix (ECM), and membranes (e.g., trans-Golgi network membrane, endosome membrane, and lysosomal membrane) (**Figure S3**). In general, we observed clear correspondence between specific VerteBrain protein clusters and well established protein complexes or biological processes, as labeled in **Table S4**.

As an example of this trend, one major central hub for protein and lipid trafficking/sorting is the golgi apparatus. The COG (<u>c</u>onserved <u>o</u>ligomeric <u>G</u>olgi) complex is an evolutionary conserved peripheral membrane protein complex that is primarily responsible for controlling membrane trafficking and ensuring Golgi homeostasis by orchestrating retrograde vesicle transport within the Golgi^45^. As expected, VerteBrain identified high confidence interactions between the eight subunits (COG1-COG8). Additionally, our data revealed interactions between COG subunits and GOLGA1 (<u>golg</u>in subfamily <u>A</u> member <u>1</u>), a member of the golgin protein family, which functions as a vesicle tethering factor to facilitate Golgi fusion events^46,47^. Specifically, VerteBrain showed that GOLGA1 interacts with COG1, COG3, and COG5, as well as GOLGA6D and COG5, COG7. To our knowledge, GOLGA1 pairwise interactions with COG1 and COG3 were previously reported in a proximity labeling study by Go *et al.*^48^ on HEK293 cells.These data suggests that the COG complex and the golgin protein family likely work together to carry out vesicle trafficking and membrane dynamics.

### VerteBrain reveals novel interactions between Rab effectors and across trafficking systems

CF-MS not only confirms known membership of protein assemblies but also reveals potential crosstalk between them. When combined with AlphaFold-Multimer modeling, this approach provides a robust framework for generating structural hypotheses and functional insights based on complex membership and interaction dynamics^31,49^. Leveraging VerteBrain’s overall accuracy, we decided to investigate such newly observed interactions among established systems.

We focused on endocytic trafficking – a fundamental cellular process essential to brain function, where it regulates key physiological activities such as synaptic signaling, receptor dynamics, and nutrient uptake^50^. Rab (<u>R</u>as-<u>a</u>ssociated <u>b</u>inding) GTPases and their effectors coordinate key steps like vesicle budding, transport, and membrane fusion^51^. One such system is the five subunit <u>e</u>ndosomal <u>R</u>ab5 and <u>R</u>NA/ribosome intermediar<u>y</u> (FERRY) complex, involved in mRNA transport on early endosomes^52^. From two *in vitro* studies^52,53^, FERRY was reported as a pentameric complex with the five subunits: TBCK (Fy1), PPP1R21 (Fy2), C12ORF4 (Fy3), CRYZL (Fy4) and GATD1 (Fy5). Our data recapitulates this assembly, suggesting that the FERRY complex forms a stable complex *in vivo*, across all species in this study (**Figure 3A & 3B**). Notably, mutations in three different members of the FERRY complex (Fy1, Fy2, and Fy3) are associated with rare neurological disorders in human patients^54^.

Similarly, VerteBrain captures high-confidence PPIs observed among subunits of the TRAPPII and TRAPPIII complexes. The TRAPP (<u>Tra</u>nsport <u>P</u>rotein <u>P</u>article) complexes are Rab GTPase exchange factors that share a core set of subunits^55^. In metazoans, there are two TRAPP complexes: TRAPPII and TRAPPIII. TRAPPII and TRAPPIII have distinct specificity for GEF activity towards Rabs, with TRAPPIII acting on Rab1, and TRAPPII acting on Rab1 and Rab11^55^. Strikingly, we also observed previously undescribed interactions between these systems. We found high CF-MS scores between TRAPPII/TRAPPIII subunits and the FERRY complex (**Figure 3B**), suggesting a network of physically and functionally connected subunits in these Rab effectors systems.

In order to further investigate interactions among Rab effectors, we specifically examined RAB5A, a key regulator of early endosomal fusion. RAB5A interacts with multiple effectors including RABEP1^56,57^ (Rabaptin, <u>Ra</u>b GTPase <u>B</u>inding <u>Ef</u>fector <u>P</u>rotein <u>1</u>) and RABGEF1 (<u>Ra</u>b <u>g</u>uanine nucleotide <u>e</u>xchange factor 1), which form a ternary complex that activates Rab5A more effectively than either protein alone^59^.

Using AlphaFold 3, we modeled this interaction and found the most favorable stoichiometry to be 2:2:2, supporting the role of RABEP1 dimerization in stabilizing the complex (**Figure 3C**). VerteBrain also recovered high-confidence interactions between RAB5A and RABGEF1, as well as between RAB5A and RABEP1. While the precise determinants of RAB5A binding remain unclear, these findings suggest that Rab5A binding is context-dependent, influenced by cellular and molecular conditions.

Given these results, we hypothesized RAB5A might also bind to multiple sites on the FERRY complex. Although direct RAB5A–FERRY interactions were not strongly supported by CF-MS scores, we found interactions between Rab effectors (RABEP1, RABGEF1) and FERRY subunits (Fy1–Fy4). To investigate these findings, we performed structural modeling using AlphaFold 3 to test the alignment of Rab5A binding with the site proposed by Quentin *et al.*^53^ (**Figure 3D**). Testing multiple configurations with increasing copies of Rab5A consistently revealed three distinct favorable binding sites, as illustrated with dotted circles labeled 1 through 3 appearing in order (**Figure 3E**). These findings highlight the dynamic and potentially promiscuous nature of Rab5A binding within the FERRY complex. Given the intrinsically disordered nature of most of the Fy2 structure, we prioritized models with the highest confidence scores. However, it is important to note that these structural models do not fully account for *in vivo* conditions, and further experimental validation is required. Further investigation is also needed to clarify the role of Rab5A in interacting with the FERRY complex and mediating endosomal processes. In the case of such context-dependent molecular assemblies the evolutionarily conserved PPIs from VerteBrain provide high quality data for hypothesis generation and testing.

### An extensive interaction network centered on Rabconnectin-3 and WDR37-PACS1/2 complexes

We examined a particular case where VerteBrain results reported previously unknown interactions between two reasonably well-conserved and defined systems; the Rabconnectin-3 complex and the WDR37-PACS1/2 complex.

Rabconnectin-3, originally identified from purified synaptic vesicle fractions in rat brain^59,60^, is comprised of Rabconnectin-3α (DMXL1 or brain-enriched DMXL2) and Rabconnectin-3β (WDR7)^61^. Despite limited characterization of individual subunit functions, DMXL1, DMXL2, and WDR7 were identified as accessory proteins specific to V-ATPase, implicating Rabconnectin-3 in synaptic vesicle acidification—a critical process in protein sorting and degradation^62^. While the original discovery of the Rabconnectin-3 complex only identified two subunits, recent studies leveraging immunoprecipitation mass spectrometry confirmed the inclusion of ROGDI^30,62^, a highly conserved metazoan protein with structural homology to yeast Rav2^63^. Since DMXL1/2 share partial homology with Rav1, another component of the yeast RAVE (<u>R</u>egulator of <u>A</u>TPase of <u>V</u>acuoles and <u>E</u>ndosomes) complex, this supports the idea that Rabconnectin-3 may function analogously to the RAVE complex as a V-ATPase chaperone^30,64^. Consistent with these observations, we constructed a 3D structural model of Rabconnectin-3 with AlphaFold 3 that shows visible similarity with the yeast RAVE complex, with the exception of WDR7 & Skp1, which as of yet have no known equivalents (**Figure 4A**).

**Figure 4.**
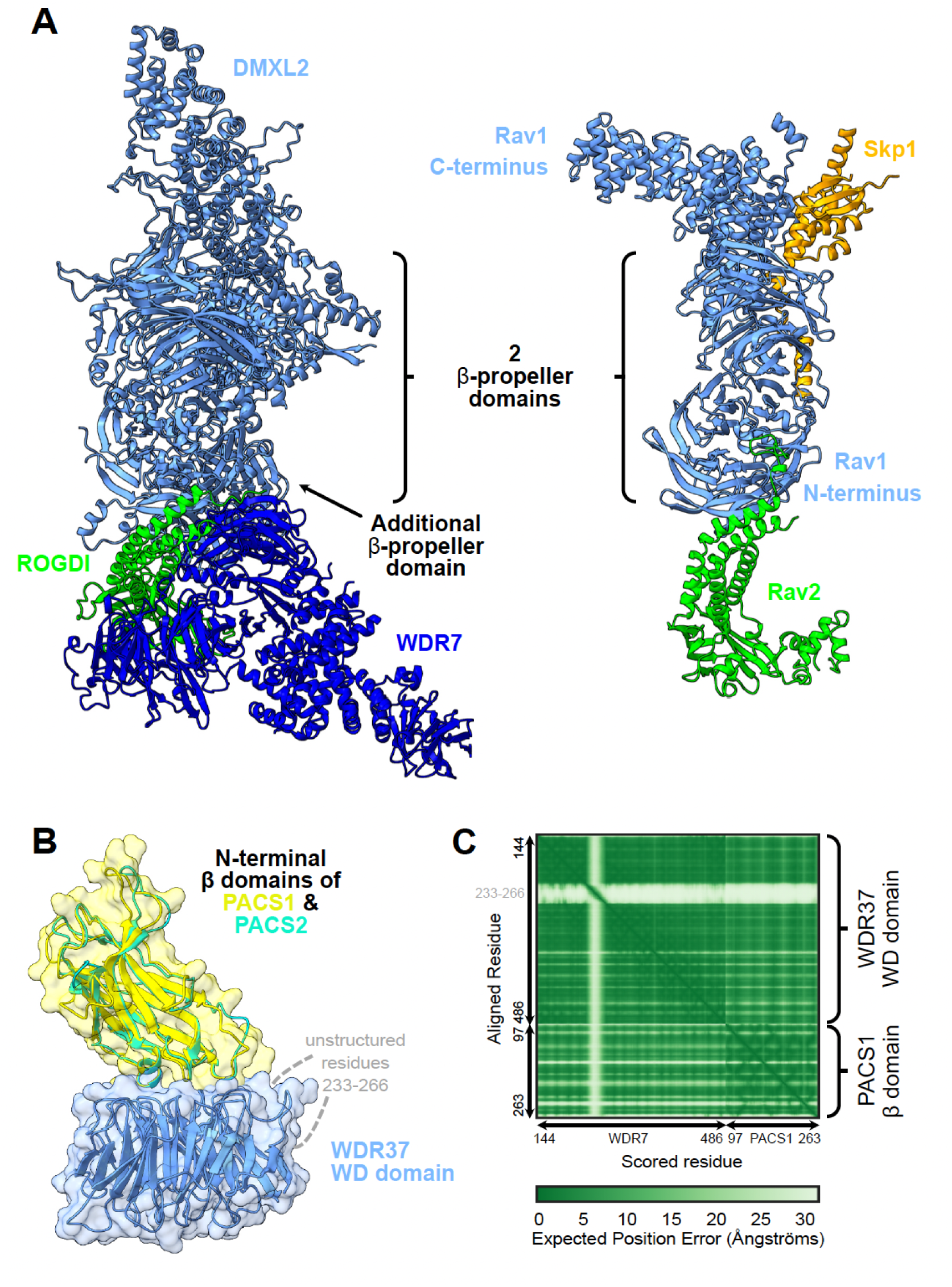
Structural predictions from AF3 reveal structural homology between the Rabconnectin-3 complex in animals and the RAVE complex in yeast. (A) AlphaFold 3 models reveal strong structural homology between the Rabconnectin-3 complex in higher eukaryotes, comprising DMXL2, WDR7, and ROGDI, and the RAVE complex in yeast, highlighting conserved architecture and potential functional parallels in V-ATPase regulation. (B) Structural modeling shows PACS1 (yellow) and PACS2 (cyan) competing for the same WDR37 binding site, suggesting dynamic, mutually exclusive interactions. (C) Predicted Aligned Error (PAE) plot showing the well-predicted WD40 domain of WDR37 interacting with the β-domain of PACS1, demonstrating high-confidence structural alignment and potential interaction stability.

Similarly, the WDR37-PACS1/2 axis is comprised of WDR37, a highly conserved WD40-repeat-containing protein with unknown function in vertebrates, and two paralogs of PACS (<u>P</u>hosphofurin <u>a</u>cidic <u>c</u>luster <u>s</u>orting) proteins, PACS1 and PACS2, which are involved in sorting cargo proteins to various organelles^65,66^. Published yeast 2-hybrid assay, co-immunoprecipitation^67^, and AP-MS^10^ experiments have confirmed the physical interaction between WDR37 and PACS1/2, although the stoichiometry remains unclear. Using AlphaFold 3, we modeled the interaction interface, which suggests that PACS1 and PACS2 compete for the same WDR37 binding site, hinting at dynamic, mutually exclusive interactions (**Figures 4B & 4C**). However, the stoichiometric and functional implications of this competition remain to be elucidated.

While VerteBrain accurately reconstructs subunit-subunit interactions for each complex, to our surprise, it also revealed previously unknown interactions between the Rabconnectin-3 (DMXL1/2, WDR7 & ROGDI) and the WDR37-PACS1/2 (WDR37, PACS1 & PACS2) axis as well as numerous shared interaction partners, suggesting the potential for functional connections between these systems. Motivated by these observations, we independently tested the interactions using immunoprecipitation-mass spectrometry (IP-MS) from pig and mouse brain lysates, targeting the Rabconnectin-3 subunits WDR7, DMXL1, DMXL2, and ROGDI, and the WDR37-PACS1/2 complex subunits WDR37 and PACS2, as well as the VerteBrain-identified interactor, RELCH, as bait proteins (**Figure 5A**). While we confirmed interactions within and between these two complexes (**Figure 5B**), we further observed a rich interaction network centered on these proteins which we visualized with a hierarchical clustering heatmap depicting the top 100 most enriched ‘prey’ proteins from our immunoprecipitation experiments (**Figure 5C** and **Table S4**). Further supporting evidence for the enrichment of these interactors is provided in **Figures S4-S7**. Notably, Rabconnectin-3 and WDR37-PACS1/2 share many common interactors, spanning elements of intracellular trafficking and signaling, and including the GABA (Gamma-aminobutyric acid) & ryanodine receptors, Na+/K+ transporting ATPases, the E3 ubiquitin-protein ligases TRIM21 & HUWE1, as well as core cellular machinery, such as the elF3 complex (**Figure 5D**).

**Figure 5.**
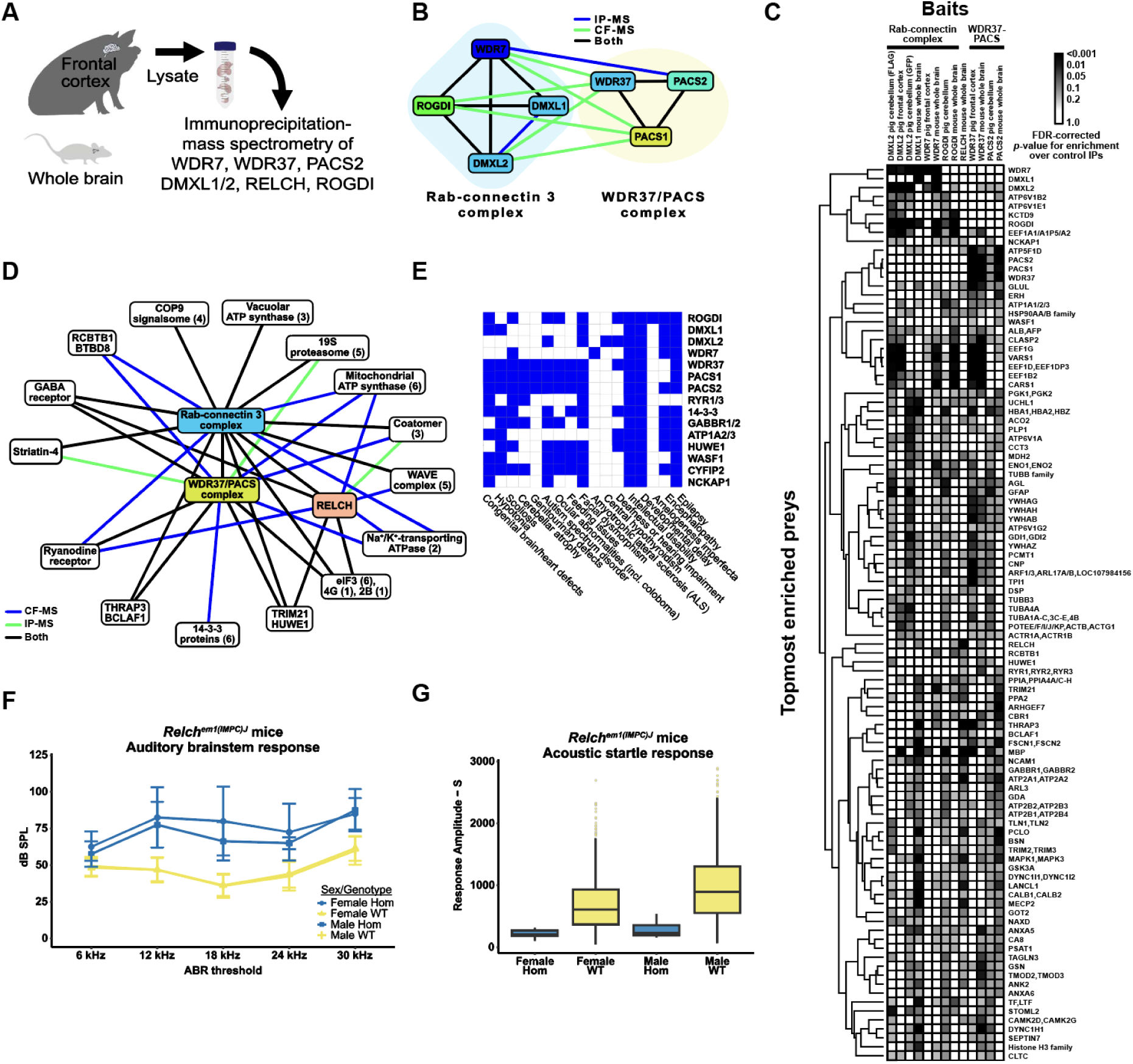
Validation of CF-MS using immunoprecipitation mass spectrometry (IP-MS) as an independent experimental method. The RAVE/Rabconnectin-3 complex interacts with WDR37/PACS1/2 in an extensive network implicated to epileptic seizures and facial dysmorphism and links the RELCH gene to deafness. (A) Schematic of the immunoprecipitation mass spectrometry (IP-MS) experiment from pig and mouse brain lysates using the indicated bait proteins (DMXL1, DMXL2, WDR7, WDR37, PACS2, ROGDI, and RELCH) to investigate protein interactions within the Rabconnectin-3 and WDR37-PACS1/2 complexes. (B) Overview of interactions between subunits of the Rabconnectin-3 and WDR37-PACS1/2 complexes, categorized by experimental evidence from IP-MS and CF-MS datasets. (C) Hierarchical clustering heatmap showing the IP-MS results for the seven bait proteins, in either pig or mouse brain protein lysates, displaying the top most enriched prey proteins. Protein interaction patterns reveal the connectivity and potential functional relationships among Rabconnectin-3, WDR37-PACS1/2, and associated complexes. (D) Summary of shared interactors identified in the Rabconnectin-3 and WDR37-PACS1/2 complexes. In most cases shown, protein interactions were detected by both IP-MS and CF-MS (complex fractionation-mass spectrometry), revealing overlapping components involved in cellular processes such as ATP synthase regulation, vesicle trafficking, and proteasome function. (E) Overview of clinical phenotypes associated with subunits of the Rabconnectin-3 and WDR37-PACS1/2 complexes. Common interactors observed in both IP-MS and CF-MS display phenotypic similarities, with disease associations including epilepsy, intellectual disability, developmental delay, and hearing impairment, highlighting potential shared pathways in neurodevelopmental disorders. (F) Auditory brainstem response (ABR) thresholds for RELCH null mice, generated by the International Mouse Consortium^79^, homozygous deletion of exons 3–5 and 454 bp of flanking intronic sequence (n=8 per sex, n=16 total), compared to healthy control mice (female n=671, male n=673). RELCH-deficient mice show significantly elevated ABR thresholds at all stimulus levels, indicating profound hearing impairment and reinforcing the role of RELCH in auditory function. (G) Acoustic startle response and pre-pulse inhibition (PPI) phenotypic assay for RELCH mutant mice with homozygous exon deletion (female n=7, male n=8, n=15 total) compared to control mice (1,949 females, 1,920 males, n=3,869 total). Mutant mice exhibit significant differences in response amplitude – decrease in startle reflex, suggesting deficits in auditory processing.

### VerteBrain reveals new connections between protein interaction and human disorders

Clinical correlations further underscore the significance of these findings. Mutations in Rabconnectin-3 components have been implicated in various neurodevelopmental disorders. For example, DMXL2 mutations and structural variants are associated with developmental delay, intellectual disability (ID), autism spectrum disorder (ASD), and epilepsy^68,69^, while ROGDI mutations cause Kohlschütter-Tönz syndrome (KTS), characterized by epilepsy, spasticity, psychomotor regression, and amelogenesis imperfecta^63,70^. Similarly, *de novo* variants in WDR37 have been linked to multisystemic syndromes characterized by various developmental anomalies, including epilepsy, developmental delay, intellectual disability, cerebellar hypoplasia, colobomas, and dysmorphic facial features^71,72^. PACS1 mutations cause intellectual disability with facial dysmorphism^73^, while PACS2 mutations are linked to developmental and epileptic encephalopathy^74^. Consistent with the protein interactions identified in VerteBrain, these disorders share overlapping clinical features, among them epilepsy, intellectual disability, and developmental delay (**Figure 5E**), reinforcing the hypothesis that Rabconnectin-3 and WDR37-PACS1/2 participate in shared biological pathways crucial for neurodevelopment.

Interestingly, previous genetic studies in humans have identified mutations in DMXL2 as pathogenic causes of dominant nonsyndromic hearing loss (<u>D</u>eafness, Autosomal Domi<u>na</u>nt 71 – DFNA71), without accompanying neurological deficits in affected patients^75,76^ (**Figure 5E**). Consistent with these findings, the zebrafish ortholog of DMXL2, localizes to the basal region of hair cells and the corresponding mutant alleles of *dmxl2* cause auditory and vestibular defects^77^. The role of DMXL2 in auditory processes was further confirmed in a recent study by Peng *et al.*^78^ demonstrating its role in endocytosis and recycling of auditory synaptic vesicles in a hair cell-specific knockout mouse model.

Building on these observations, our VerteBrain analysis reports a perfect pairwise score of 1 between RELCH (<u>R</u>ab11-binding and <u>L</u>isH domain, <u>c</u>oiled-coil and <u>H</u>EAT repeat-containing) and DMXL2, suggesting a strong functional or physical association. In IP-MS experiments using Rabconnectin-3 subunits as bait proteins, RELCH emerged as a high-confidence interactor (**Figure 5D**). Likewise, homozygous RELCH mutant mice displayed abnormal auditory responses (**Figure 5F**) and a reduced startle reflex (**Figure 5G**), as reported by the International Mouse Genetic Consortium^79^ (see **Methods**). Taken together, these findings drawn from VeterBrain observations as well as IP-MS validation bridge evidence from human genetics, zebrafish models, and mouse studies, to collectively point to RELCH as a potential novel causative gene for deafness in humans.

Other noteworthy interactors of the Rabconnectin-3 and WDR37/PACS1-2 complexes identified in VerteBrain include the <u>ga</u>mma-amino<u>b</u>utyric <u>a</u>cid (GABA) receptors, the major inhibitory neurotransmitter receptors in vertebrate brain. Specifically, we observed both complexes to interact with GABA receptor subunits, both by CF-MS and IP-MS experiments, an observation replicated with 4 independent IP-MS baits (ROGDI, DMXL1, RELCH, and PACS2). Mutations in different subunits have been associated with genetic epilepsy syndromes with different clinical phenotypes^80,81^.

Our identification of RELCH and GABA receptors as shared interactors within these conserved complexes provides a new framework for studying their roles in human disorders. These findings expand our understanding of intracellular trafficking networks and lay the groundwork for future research into their implications for epilepsy, intellectual disability, and hearing impairment. While many of the clinical features described here are common across diverse, unrelated neurodevelopmental disorders, the involvement of a shared cellular mechanism between these proteins underscores the importance of exploring a potential convergent pathological pathway.

### Conserved interactions extending beyond the brain: INPPL1, ARHGEF1, and short stature disorders

Patients with neurological deficits, such as ASD, ID, and developmental delay, often exhibit additional syndromic anomalies, including skeletal defects like scoliosis or hypotonia^82,83^ (**Figure 5E**). Although these clinical manifestations are seemingly distinct, growing evidence highlights the shared molecular pathways and cellular mechanisms underpinning both brain and skeletal development. PPI networks in the brain can reveal critical regulatory nodes that extend their influence beyond neural phenotypes, shedding light on systemic processes like cytoskeletal organization, cellular signaling, and tissue homeostasis. Notably, skeletal dysplasias are increasingly recognized to co-occur with neurological symptoms^84,85^, suggesting that disruptions in shared developmental pathways can simultaneously impact both the brain and the skeleton. Motivated by this broader perspective, we turned our focus to INPPL1 (<u>in</u>ositol <u>p</u>olyphosphate <u>p</u>hosphatase-like 1), a gene with reported mutations linked to the rare human endochondral bone disorder, opsismodysplasia (OPS), that affects growth plate expansion and skeletal development^86–88^. Clinical features of OPS include short stature with short limbs, small hands and feet; macrocephaly with a prominent anterior fontanel; and facial dysmorphism, although severity varies at birth.

The *INPPL1* gene encodes the 5’-inositol phosphatase SHIP2 (<u>s</u>rc <u>h</u>omology 2 domain-containing inositol <u>p</u>hosphatase 2), which regulates phosphoinositide signaling by removing the 5’-phosphate from substrates such as PI_(3,4,5)_P_3_ and PI_(4,5)_P_2_ in cellular membranes^89,90^. Previous work by Voigt *et al*.^91^ demonstrated that homozygous mutations in *inppl1a*, the zebrafish ortholog of INPPL1, cause defects in notochord vacuolated cell expansion and hypertrophic chondrocyte differentiation leading to vertebral malformations and scoliosis and endochondral bone defects in zebrafish. VerteBrain revealed several high CF-MS scoring protein interactions with INPPL1. We filtered these candidate interactors using AlphaFold 3 to test for putative direct binding, and found a medium level of confidence for direct binding with ARHGEF1 (<u>Rh</u>o guanine nucleotide exchange factor 1) (**Figure 6A-C**). ARHGEF1 is an intracellular signaling molecule that has been shown to modulate signaling from G-protein-coupled receptors to RhoA, and as such, it is an important regulator of the actin cytoskeleton with pleiotropic functions^92^. Interestingly, ARHGEF1 is primarily implicated in immunological functions, as its loss causes a primary antibody deficiency in humans^93^. This observation highlights the heterogeneity of disease etiology, as disruption of a single gene can affect multiple pathways, resulting in diverse clinical manifestations.

**Figure 6.**
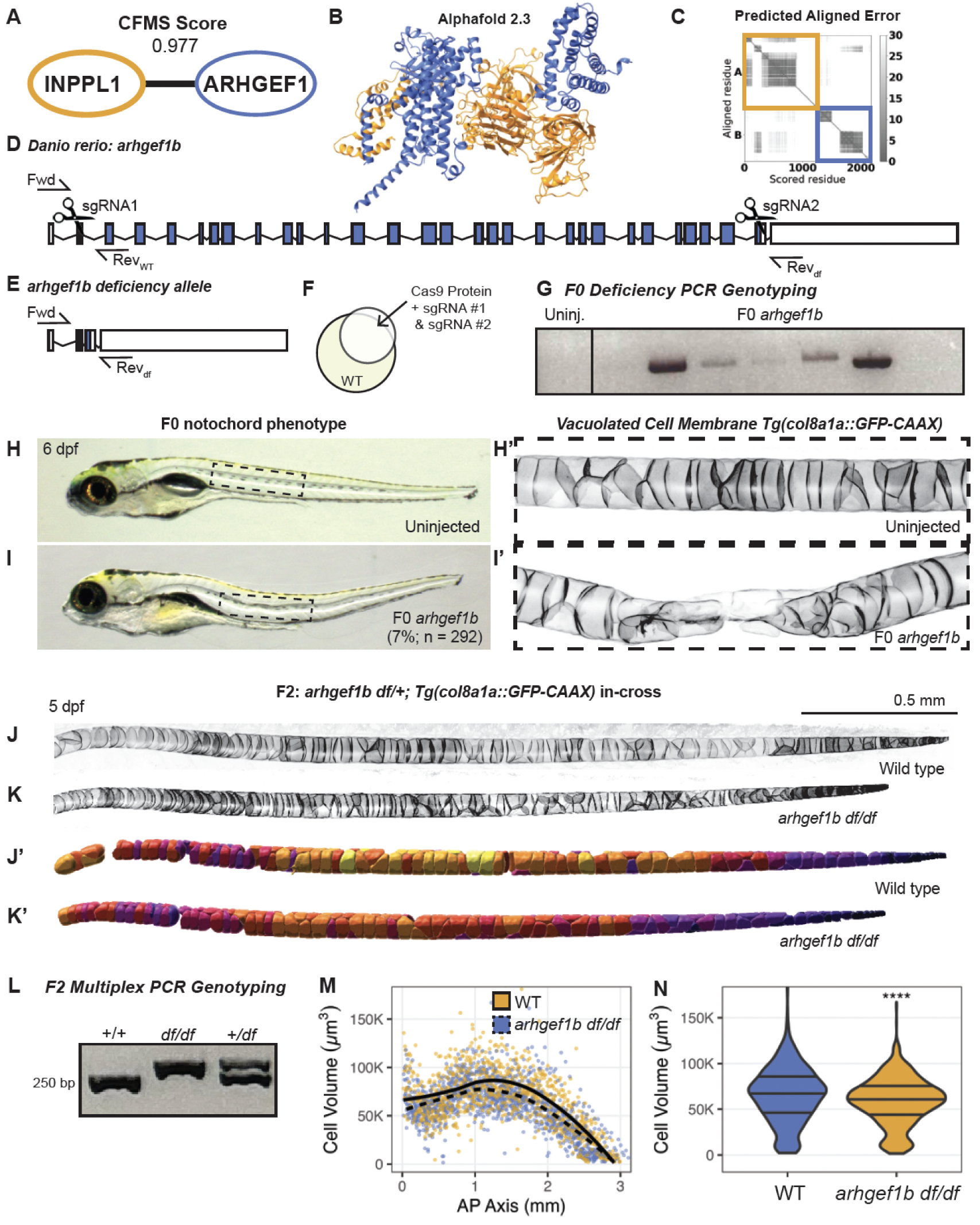
VerteBrain identifies novel interactors of INPPL1 important for skeletal development and disorders. (A) INPPL1 BrainMAP cluster identified an interaction between INPPL1 and ARHGEF1, with a CF-MS score of 0.977. (B) Structural modeling of INPPL1-ARHGEF1 interaction using AlphaFold 2.3 Multimer. INPPL1 is depicted in gold, while ARHGEF1 is depicted in blue. (C) Predicted aligned error (PAE) plot of INPPL1-ARHGEF1 interaction. Gold box indicates INPPL1 residues, while the blue box indicates ARHGEF1 residues. (D) Genomic structure of *D. rerio arhgef1b* locus (White boxes represent untranslated exonic regions, blue boxes represent translated exonic regions, lines represent intronic regions) with CRISPR mutagenesis strategy (scissors represent CRISPR sgRNA targeting loci, arrows represent approximate locations of the various genotyping primers: Fwd = universal forward primer, Rev-WT = reverse primer to detect the wild-type allele, Rev-df = reverse primer to detect the deficiency allele). (E) Genomic structure of the *D. rerio arhgef1b* deficiency allele construct with alignment of relevant genotyping primers (Fwd = universal forward primer, Rev-df = reverse primer to detect the deficiency allele). (F) CRISPR mutagenesis was achieved via 1 nL injection of Cas9 protein, sgRNA1 targeting the *arhgef1b* start site, and sgRNA2 targeting the *arhgef1b* stop site of the locus into one-cell staged wild-type zebrafish embryos. (G) PCR validation in 1 dpf F0 *arhgef1b* crispant embryos demonstrating that the *arhgef1b* deficiency allele was generated by the CRISPR mutagenesis strategy. Uninjected (Uninj.) controls amplicons are absent when using the universal forward primer and the deficiency-specific reverse primer, while pooled F0 crispants display amplicons. (H) Representative lateral view of uninjected control embryos at 6 dpf, with (H’) inset showing max projection of confocal images of the notochord vacuolated cell membrane marker: *Tg(col8a1a::GFP-CAAX)*. (I) Representative lateral view of F0 *arhgef1b* crispant embryos at 6 dpf showing body curvatures (7%, n = 292), with (I’) inset showing max projection of confocal images of the notochord vacuolated cell membrane marker: *Tg(col8a1a::GFP-CAAX)*, showing vacuolated cell collapse at region of body curvature. (J) Representative max projections of confocal stacks of notochord vacuolated cell membranes labeled by the *Tg(col8a1a::GFP-Caax)* transgene in 5 dpf wild-type progeny of *arhgef1b df/+* in-cross, with (J’) IMARIS Cell Segmentation of colored by cell volume (Scale bar = 0.5 mm). (K) Representative max projections of confocal stacks of notochord vacuolated cell membranes labeled by the *Tg(col8a1a::GFP-Caax)* transgene in 5 dpf homozygous mutant progeny of *arhgef1b df/+* in-cross, with (K’) IMARIS Cell Segmentation of colored by cell volume (Scale bar = 0.5). (L) Representative agarose gel of the duplex PCR genotyping method used to distinguish wild-type (+/+), homozygous mutant (df/df) and heterozygous carrier (df/+) larvae used in imaging experiments. (M) Scatterplot of cell volumes (μm^3^) plotted against their anterior-posterior (AP) axis (mm) at 5 dpf in wild type (orange color palette, n = 922 cells, N = 7 fish) and *arhgef1b df/df* homozygous mutants (blue color palette, n = 1318 cells, N = 10 fish) from *arhgef1b df/+* in-crosses, with loess lines of best fit for each genotype. (N) Violin plot of cell volumes (μm^3^) of 5 dpf in wild type (orange color palette, n = 922 cells, N = 7 fish) and *arhgef1b df/df* homozygous mutants (blue color palette, n = 1318 cells, N = 10 fish) from *arhgef1b df/+* in-crosses (Horizontal lines represent 25th, 50th, and 75th quartiles, **** indicates p-value less than 0.0001).

To further explore the functional relationship between INPPL1 and ARHGEF1, we generated a null allele of *ARHGEF1* in zebrafish. Although zebrafish have two paralogs of *ARHGEF1* (*arhgef1a* and *arhgef1b)* RNA-sequencing data from Daniocell indicate *arhgef1a* is primarily expressed in hematopoietic cells^94^, similar to reported expression patterns in mouse^95^. In contrast, *arhgef1b* is more broadly expressed in developing zebrafish, suggesting that this paralog functions in extra-hematopoietic systems. To probe these potential roles of *arhgef1b*, we generated a null allele using a CRISPR method with small guide RNAs (sgRNAs) targeting the 5’– and 3’-ends of the gene (**Figure 6D-F**, see **Methods**). To better visualize potential notochord defects, we performed the CRISPR injections in a transgenic line expressing membrane-localized GFP in notochord vacuolated cells^95^. This approach successfully generated what will be referred to as the *arhgef1b* deficiency (*df*) allele, as much of the coding region is deleted and confirmed by PCR (**Figure 6D,E,G**).

At 6 days post fertilization (dpf), 7% of the genetically mosaic F0 *arhgef1b* crispants displayed body curvatures and notochord defects (**Figure 6H-I**; n = 292), and confocal microscopy revealed collapse of notochord vacuolated cells at the sites of body curvature (**Figure 6I’**), supporting its role in vacuolated cell biogenesis. F1 *arhgef1b^df/+^* heterozygotes were in-crossed to assess vacuolated cell and notochord morphology in *arhgef1b^df/df^* homozygous mutants (**Figure 6J-K**). Analysis of vacuolated cell volumes using a semi-automated cell segmentation (IMARIS) (**Figure J’-K’)** demonstrated a significant decrease in vacuolated cell volume in *arhgef1b^df/df^* mutants compared to wild-type siblings at 5 dpf (**Figure 6M-N**), which were genotyped using a multiplex PCR strategy to identify zygosity (**Figure 6L**, see **Methods**).

Homozygous *arhgef1b^df/df^* mutants did not survive to adulthood, and we were therefore unable to assess skeletal phenotypes in *arhgef1b^df/df^* adults. We suspect that the broad expression of *arhgef1b* indicates a role in a wide variety of tissues, any of which could be essential for survival. Regardless, these embryonic results strongly support a role for *arhgef1b* in vacuolated cell size regulation, similar to that of *inppl1a*, and suggest a role for INPPL1-ARHGEF1 interactions in notochord development in zebrafish. Notably, a protein interaction observed in the brain successfully predicted a phenotype in a second tissue, indicating that many brain interactions may be relevant to multiple tissue types.

## CONCLUSIONS

Here, we present a compendium of protein-protein interactions conserved across vertebrate brains, spanning over 300 million years of evolution. VerteBrain serves as a valuable resource for uncovering conserved molecular assemblies in vertebrate brains. We demonstrate how these interactions can be applied to discover human disease-associated protein functions, such as the link between INPPL1-ARHGEF1 and short stature disorders, as validated in zebrafish, and complex synaptic vesicle trafficking modules associated with epilepsy and congenital deafness. These findings highlight the potential of VerteBrain as a resource for identifying mechanisms underlying neurological disorders and guiding future therapeutic research.

## AUTHOR CONTRIBUTIONS

Design and co-supervision: V.D. and E.M.M.

Proteomics samples and experiments: V.D., aided by G.H., O.P., R.P., J.C.L., and B.A.N.

Data analysis: V.D., aided by D.Y., M.L., R.M.C and C.D.M.

Zebrafish experiments, design, and supervision: B.V. and R.S.G.

Manuscript initial draft: V.D, B.V, and E.M.M.

All authors discussed results and contributed edits.

## MATERIALS AND METHODS

### Sources and handling of brain samples

A Dolphin (*Tursiops erebennus*) brain was obtained from a juvenile 180 cm female *Tursiops erebennus* (Individual # MMES2016042SC; SC1629; 6-25-16) according to established necropsy standard operation procedures at the NOAA/NCCOS laboratory in Charleston, South Carolina under federal employee authority as outlined in Section 109(h) of the Marine Mammal Protection Act. The animal was positive for *Brucella ceti* in the lung and brain. The brain sample was collected and stored at –80 C, prior to shipping on dry ice for proteomic analysis under NOAA/NMFS permit 24359 as an Authorized Recipient (June 8, 2023). Because this dolphin species was only recently recognized as a coastal ecotype of the common bottlenose dolphin (*Tursiops truncatus*), and a reference proteome for *T. erebennus* is not yet available, we used the genome of *T. truncatus* (common bottlenose dolphin) for all computational analyses.

Frozen brains were obtained from the following additional vertebrate species as follows: 5 whole pig brains (Sus scrofa), 20 whole chicken brains (*Gallus gallus*) and 50 whole mouse brains (Mus musculus) from animals of unknown age/sex were obtained from Sierra for Medical Sciences (Whittier, California). 20 frozen rabbit (*Oryctolagus cuniculus*) brains were obtained from Pel-Freez Biologicals (Rogers, Arkansas). These brains were all harvested and immediately flash frozen prior to shipping on dry ice. Dissection of specific brain regions was performed in house as described below. 35 whole zebrafish (*Danio rerio, Tubingen strain,* 8-10 months post fertilization) brains were harvested in house and immediately flash frozen (see methods below). Both male and female zebrafish were used in approximately equal numbers.

### Sample preparation, protein extraction, and co-fractionation/mass spectrometry

#### Dissection of pig and rabbit brains

Unless stated otherwise, all steps were performed on ice, and prior to dissection, we removed the pia mater of the brains using surgical design Teflon thumb forceps (Fisher Scientific) and washed gently with PBS pH 7.4 (Gibco, Thermo Scientific #10010023).

Pig brains were dissected immediately after receiving from the supplier using a disposable scalpel with a carbon steel blade into the following regions: frontal cortex, temporal lobe, occipital lobe, and cerebellum. Rabbit brains were thawed on ice until tissues were soft enough to dissect (regions as above). Following dissection, brain regions were flash frozen in liquid nitrogen and stored in –80°C until the proteomics analysis.

#### Native protein extraction

Unless stated otherwise, all samples were ground to a fine powder in liquid nitrogen using a chilled mortar and pestle. Powder was resuspended in an approximately equal volume of ice cold lysis buffer (HEPES lysis buffer with NP-40, 2X, Thermo Scientific), with the addition of an EDTA-free protease inhibitor cocktail tablet (cOmplete, Roche) and phosphatase inhibitor tablet (PhosSTOP, Roche). The mixture was homogenized on ice using a chilled 2mL glass dounce with approximately 20-30 strokes by hand and incubated on ice for 30 minutes. The following steps were then conducted at 4°C: The crude homogenate was clarified twice using a tabletop centrifuge (5430R, Eppendorf) at 15,000 for 20 minutes. Further clarification of the supernatant was performed by ultracentrifugation at 30,000 x g for 1.25 hours in an NVT65.2 rotor (Beckman Coulter).

#### Zebrafish brain dissection and protein extraction

Sexually mature *Danio rerio* (8-10 mpf, Tubingen strain) were euthanized in chilled 0.4% MS-222 (Tricaine, C_10_H_15_NO_5_S) before being transferred to chilled phosphate-buffered saline (PBS). The operculum and jaw bones were removed with sharp forceps. The parietal and frontal bones of the skull were chipped away to expose the brain, which was removed by severing the spinal cord posterior to the medulla. The surrounding tissue was pulled away, and the eyes were removed by severing the optic nerves. Once isolated, the dissected brain tissue was briefly rinsed in PBS, transferred to a Protein LoBind tube (Eppendorf #022431081), and snap frozen in dry ice. Samples were stored at –80 °C prior to lysis. For each chromatographic separation, brain tissue was pooled from 15-20 individuals in an approximately equal volume of chilled Pierce IP Lysis Buffer (Thermo Scientific #87788) with freshly added protease (cOmplete Mini, EDTA-free; #11836170001) and phosphatase (Roche PhosSTOP; #04906837001) inhibitors. The suspension was homogenized with a motorized teflon pestle (60 passes) and in a 2 mL glass dounce (15 passes of pestle A then 15 passes of pestle B). The homogenized lysate was incubated on ice for 30 minutes with extensive pipetting every 10 minutes before centrifugation at 15,000 x *g* for 15 minutes at 4 °C. The supernatant was transferred to a pre-chilled LoBind Protein tube, snap-frozen in liquid nitrogen or dry ice, and stored at –80 °C until further processing. Further clarification of the supernatant was performed by ultracentrifugation at 30,000 x g for 1.25 hours in an NVT65.2 rotor (Beckman Coulter) directly prior to use.

#### HPLC chromatography

Brain lysates were fractionated on a Dionex UltiMate3000 HPLC system consisting of an RS pump, Diode Array Detector, and Conductivity Monitor, Automated Fraction Collector (Thermo Scientific, CA, USA) and a Rheodyne MXII Valve (IDEX Health & Science LLC, Rohnert Park, CA) using biocompatible PEEK tubing and either size exclusion chromatography or one of two ion exchange separations (mixed bed, or triple-phase WAXWAXCAT, see below). We loaded 1-4 mg protein per separation as measured by DC Protein Assay (BioRad), collecting chromatography fractions in 2mL 96-deep well plates (Eppendorf). Select support ribs in the base were notched with a single-edged razor blade prior to fraction collection to accommodate subsequent use of the Life Technologies magnetic plate for on-bead tryptic digest during mass spectrometry sample preparation^96^.

#### Size exclusion chromatography (SEC)

BioSep-SEC-s4000 600 3 7.8 mm ID, particle diameter 5 mm, pore diameter 500 A ° (Phenomenex, Torrance, CA). Prior to chromatography of brain lysates, molecular weight standards (Sigma –Aldrich, MWGF1000, 2-5 ug each of Thyroglobulin (T9145), b-amylase (A8781), and bovine serum albumin (A8531)) were fractionated for molecular weight estimation. Each chromatography run was conducted with mobile phase PBS pH 7.4 (Gibco), 200µL load volume, flow rate 0.5mL/min, and fraction collection every 45 seconds (0.375 mL).

#### Mixed bed ion exchange chromatography (IEX)

Poly CATWAX A (PolyLC Mixed-Bed WAX-WCX) 200 3 4.6 mm ID, Particle diameter 5 mm, pore diameter 100 A ° (PolyLC Inc., Columbia, MD) comprises cation-exchange (PolyCAT A) and anion-exchange (PolyWAX LP) materials in equal amounts. Each chromatography run was loaded with 250µL sample at ⪯ 40 mM NaCl. Fractionation was performed with a flow rate of 0.5 mL/min, fraction collection every 45 seconds (0.375 mL) and a 60-minute gradient elution of 0-70% Buffer B: Buffer A (10 mM Tris-HCl pH 7.5, 5% glycerol, 0.01% NaN3), Buffer B (1.5 M NaCl in Buffer A).

#### Triple phase ion exchange chromatography (WWC)

Three columns, each 200 3 4.6 mm ID, particle diameter 5 mm, pore diameter 100 A °, were connected in series in the following order: two PolyWAX LP columns followed by a single PolyCAT A (PolyLC, Inc, Columbia, MD). Loading, buffers, and fraction collection were as for mixed bed ion exchange above with slight modifications in flow rate and elution from the methods of Havugimana *et al.* (2012). The flow rate was 0.25 ml/min with a 120-minute gradient from 5%–100% Buffer B.

#### Mass spectrometry sample preparation

Samples were prepared for mass spectrometry in 96-well plate format using either Method 1 (ultrafiltration and in-solution digest) or Method 2 (on-bead digest) MS preparation protocols from McWhite *et al.*, 2021^96^ (STAR Protocols).

#### Mass spectrometry data acquisition and processing

*Acquisition*. Mass spectra were acquired using one of two Thermo mass spectrometers: Orbitrap Fusion or Orbitrap Fusion Lumos. In all cases, peptides were separated using reverse phase chromatography on a Dionex Ultimate 3000 RSLCnano UHPLC system (Thermo Scientific) with a C18 trap to C18 column configuration. Peptides were eluted using a 3%–45% acetonitrile gradient in 0.1% formic acid over 60 min. For the Orbitrap Fusion chromatography was on an Acclaim C18 PepMap RSLC column (Dionex; Thermo Scientific) with a nano-electrospray source. For the Orbitrap Fusion Lumos both chromatography and electrospray were achieved with an EASY-Spray PepMap RSLC C18 column (Thermo Scientific, ES902).

#### Orbitrap Fusion

Top speed CID with full precursor ion scans (MS1) collected at 120,000 m/z resolution and a cycle time of 3 sec. Monoisotopic pre-cursor selection and charge-state screening were enabled, with ions of charge > + 1 selected for collision-induced dissociation (CID). Dynamic exclusion was active with 60 s exclusion for ions selected once within a 60 s window. For some experiments, a similar top speed method was used with dynamic exclusion of 30 s for ions selected once within a 30s window and high energy-induced dissociation (HCD) collision energy 31% stepped +/4%. All MS2 scans were centroid and done in rapid mode.

#### Orbitrap Lumos

Top speed HCD with full precursor ion scans (MS1) collected at 120,000 m/z resolution. Monoisotopic precursor selection and charge-state screening were enabled using Advanced Peak Determination (APD), with ions of charge > + 1 selected for high energy-induced dissociation (HCD) with collision energy 30% stepped +/ 3%. Dynamic exclusion was active with 20 s exclusion for ions selected twice within a 20s window. All MS2 scans were centroid and done in rapid mode.

### Computational analyses of protein mass spectrometry datasets

#### Reference proteome databases

Reference proteomes for *Sus scrofa* (Taxon ID: 9823, UP000008227), *Gallus gallus* (Taxon ID: 9031, UP000000539), *Oryctolagus cuniculus (*Taxon ID: 9986, UP000001811), *Mus musculus (*Taxon ID: 10090, UP000000589) *, Tursiops truncatus* (Taxon ID: 9739, UP000245320) *Danio rerio* (Taxon ID: 7955, UP000000437) and *Homo sapiens* (Taxon ID: 9606, UP000005640) were downloaded from https://www.uniprot.org/ (The UniProt Consortium, 2019) in August 2020.

#### Orthogroup assignment & curation of orthogroup-based protein reference databases

Each reference proteome protein was assigned to an eggNOG vertebrata (verNOG, taxonomic level = 40674) orthogroup using the eggNOG-mapper v5 software and the eggNOG v5.0 diamond mapper (Huerta-Cepas *et al.*, 2019^18^). Each set of proteins across all species assigned to the same orthogroup were operationally considered to belong to the same orthogroup for subsequent analyses. We updated each mass spectrometry reference proteome database by concatenating protein sequences from the same orthogroup into single entries with triple-lysines separating each protein, thus computing proteomics support on an orthogroup-by-orthgroup basis, rather than a protein-by-protein basis, which allowed for direct comparisons of proteomics data across the full vertebrate dataset.

#### Initial assignment of peptide mass spectra

For downstream analysis Thermo Fisher raw files were converted into mzXML format using GUI MSConvert^97^. Peptide inference was performed with MSGF+, X!Tandem, and Comet-2013020, each run with 10ppm precursor tolerance, and allowing for fixed cysteine carbamidomethylation (+57.021464) and optional methionine oxidation (+15.9949). Peptide search results were integrated with MSBlender (Kwon et al., 2011^19^), available from https://github.com/marcottelab/run_msblender.

For a given sample and mass spectrometry experiment, peptide spectral matches (PSMs) were analyzed as follows: for each reference database orthogroup, we summed the PSMs that could be uniquely attributed to that orthogroup. In this way, we avoided double-counting PSMs across highly related paralogs and thus near duplicate proteins, and we did not otherwise have to consider variable numbers of proteins or their relative lengths within each orthogroup in a species.

### Identification and scoring of protein-protein interactions

#### Assembly of features for scoring putative protein interactions

For each experiment, we assembled an elution matrix of identified proteins with verNOG ID (rows) by fractions (columns). Additionally, we concatenated the elution matrices across all experiments within each species as four additional matrices. Next, we calculated a series of all-by-all pairwise scores between orthogroups for individual matrices and the concatenated matrices.

We focused our analysis on well-observed proteins, so we additionally filtered for proteins with at least 60 total PSMs observed across the 30 combined fractionations. The scores/features were as follows: (1) Pearson’s r, (2) Spearman’s rho, (3) Euclidean distance, (4) Bray-Curtis similarity, (Drew *et al*., 2017)^98^. All features/scores were calculated with added Poisson noise as in Drew *et al*.^98^. Euclidean distance and Bray-Curtis similarity scores were inverted and normalized to a max score of 1. For each individual matrix from each fractionation, features 1–4 were calculated using the extract_features.py script from https://github.com/marcottelab/protein_complex_maps/

#### Construction of the gold standard protein complex training and test sets

We used known mammalian protein complexes (human, mouse and rat) from the CORUM 4.1^27^ release database (available at https://mips.helmholtz-muenchen.de/corum/download) as a gold standard positive set of stable protein-protein interactions. Known complexes were divided into positive training and test complexes according to the scheme from Cox *et al.*^17^ (2024) and complexes with over 30 members removed. To address class imbalance, the total number of negative labels was restricted to 3X the observed number of positive PPIs in our data, resulting in 3,123 total positive PPIs and 9,369 total negative PPIs labels in our feature matrix. Note that for computing training/test sets and for identification and scoring of protein-protein interactions, we used the software pipeline developed for the LECA protein interactome^17^, available in full at https://github.com/marcottelab/leca-proteomics.

Positive PPIs that participate in multiple complexes are a potential source of representation bias in the truth set, which can lead to under or overfitting during model training. To overcome this, we implemented a data stratification approach, as described in Cox *et al.*^17^

#### Identification of interacting proteins by supervised machine learning

We utilized TPOT to train our machine learning model to find pairwise interactions among members of protein complexes. We first used the scikit-learn ExtraTreesClassifier feature selection to reduce the dimensionality of the feature matrix to the top 100 features based on declining feature importance (see Zenodo repository for data). We used the TPOT (Olson and Moore, 2016) AutoML wrapper of scikit-learn machine learning functions for all subsequent training steps. A group-based split method (scikit-learn’s “GroupShuffleSplit” class) was used to generate 5 sets of test and training data, where approximately 75% of the labeled data was used for training and 25% for testing. Each of the 5 training sets were used as input into TPOT (Olson and Moore, 2016) an automated machine learning pipeline built on top of scikit-learn, to find the best classification method, pre-processing steps, and parameters for our data.

#### Construction of the protein interaction network

Interaction scores above a 8% false discovery rate threshold (CF-MS score >= 0.88) were used to build a PPI-based orthogroup network. In the network, nodes are verNOG ID mapped for corresponding proteins. This resulted in a network with 6,108 nodes and 81,225 edges. Two nodes (orthogroups) connected by an edge in this network indicate at least one interacting protein pair between the two nodes. Cuts closer to the root of the dendrogram result in larger and less granular complexes and cut closer to the tips defined smaller and finer granular subcomplexes.

### Independent validation of CF-MS interactions by comparison to external protein interaction datasets

We classified the external datasets into two categories: all-by-all and bait-prey. For all-by-all (HuRI, mouse synaptosome, hu.MAP 2.0) we treated all 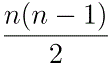 possible protein pairs as tested interactions. For bait-prey (AP-MS, OpenCell) we treated all possible bait-prey pairs as tested interactions, resulting in a total of 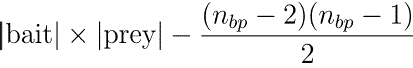 possible interactions where 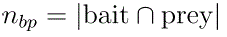. Any possible protein pair that was not classified as a positive protein pair was then classified as a negative protein pair. We divided the score range into 10 bins, 10 bins 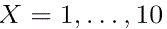, such that the coverage of each bin was 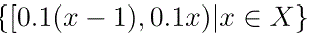, with the exception of the final bin, which covered the range 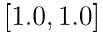. This final bin was included to capture the highest possible score from the classifier.

For each bin, we performed the following:

> We obtained a set of protein pairs from CF-MS for the given bin. This was the set of positive CF-MS protein pairs for a bin CFMS^+^. We extracted all individual proteins CFMS_single_ from the protein pairs CFMS^+^ and enumerated a set of all possible CF-MS pairs CFMS_all_ using all-by-all (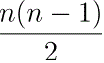 pairs). For a given bin, we ended up with 2 protein pair sets – CFMS^+^ and CFMS_all_.

We then repeated the following for each external data source *E* and for each bin:

> We obtained the set of individual proteins from the protein pairs from CF-MS for a given bin CFMS_single_. We retrieved the set of individual proteins for *E* (this does not change for a given bin since the external data’s PPI value is binary rather than score-based), which we denoted as *E*_single_. Then, we calculated the intersection between the two sets of individual proteins 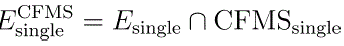. From 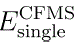, depending on the methodology of all-by-all or bait-prey as described earlier, we obtained our set of all possible protein pairs *E*_all_. We calculated another intersection between *E*_all_ and the set of protein pairs from *E* to determine the set of positive *E* protein pairs for a specific bin, which we denoted as *E*^+^. The negative pairs were determined by 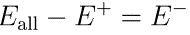
>
> In all, we determined the following sets for a given bin and external dataset of protein pairs *E*:
>
> CFMS^+^ – Set of positive CFMS protein pairs
>
> CFMS_single_ – Set of all observed individual CFMS proteins
>
> CFMS_all_ – Set of all possible CFMS protein pairs
>
> *E*_single_ – Set of all observed individual external data proteins
>
> 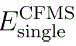 – Set of positive external data proteins *E*_single_ that are also found in CFMS_single_
>
> *E*_all_ – Set of all possible external data proteins from 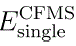
>
> *E*^+^ – Set of all positive external protein pairs in *E*_all_ i.e. protein pairs found in both *E* and *E*_all_
>
> *E*^−^ – Set of all negative external protein pairs in *E*_all_ i.e. protein pairs in *E*_all_ that are not found in *E*
>
> The enrichment score was calculated as 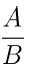 where 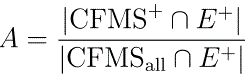 and 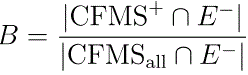. The enrichment score is thus an indicator of how well the set of CFMS positive PPI overlaps with the set of external positive PPI while minimizing the overlap between CFMS positive PPI and external negative PPI for a given bin.

### Evaluating pairwise interactions for evidence of direct binding using AlphaFold-Multimer

We analyzed all candidate pairs using ColabFold 2.3, which uses the same architecture and weights as AlphaFold 2.3. We evaluated all ColabFold results for direct binding by observing 4 scores – ipTM (interface predicted TM-score), iPAE (interface predicted aligned error) 10th percentile, iPAE minimum, and pDockQ (predicted DockQ). The ipTM is produced directly from AlphaFold-Multimer. The PAE is produced directly from AlphaFold and represents an *n × n* matrix where *n* represents the combined length of the protein pair. We derived the iPAE by taking the inter-chain PAE values. We used a cutoff of pDockQ ≥ 0.23 for an acceptable direct interaction between two proteins, as was used in the original pDockQ and DockQ papers.

### Evaluating the predictive power of the measured interactions for biological processes

To measure the prediction power of our network, we tested whether we could predict novel proteins related to specific biological annotations (GO biological processes) given already known human proteins (seed sets) for the annotation.

We made a list of human proteins for each annotation and mapped these proteins to vertebrate-level orthogroups. We filtered out orthogroups not belonging to the network. Annotations having at least five orthogroups (in the network) were used for the following steps. We ranked all orthogroups in the network for each annotation based on the sum of edge weights linked to the seed set. We annotated a new score *S_i_* for each orthogroup *i* following this formula:

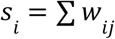

Where *j* is an orthogroup in the seed set, and *w*_*ij*_ is the weight of the edge connecting *i* and *j*. Then orthogroups were ranked by their *s*_*i*_. (There is no self-link, i.e., *i* ≠ *j*)

Then, we measure the predictability of the network for each annotation by calculating the area under the receiver operating characteristic curve (AUROC). This value ranges from 0 to 1. 0.5 means a random classifier, and 1 represents a perfect classifier. The true positive rate (TPR) and false positive rate (FPR) were calculated by this formula:

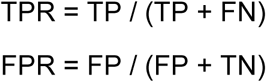

Where TP (true positive) is the number of orthogroups belong the seed set and ranked above a threshold, FP (false positive) is the number of orthogroups not in the seed set but ranked above a threshold, FN (false negative) is the number of orthogroup in seed set and ranked below a threshold, and TN (true negative) is the number of orthogroups not in seed set and ranked below a threshold. We also calculated AUROC values with random seed sets, which have the same sizes as the annotations. Then, we compared the distribution of AUROC values for annotations of interest and the random.

### Mapping of orthogroups to Disease Ontology terms

Human disease-gene associations data were retrieved from the Alliance of Genome Resource (Ver 5.3.0, Oct 28, 2022). Among five different types of associations, only “Is Implicated In” genes were used, so genes whose variants are known to function in causing or modifying diseases were assigned. Then, vertebrate-level orthogroups were mapped to each disease term based on the assigned human genes.

We inherited orthogroups of ancestor terms to their child terms following the structure of Disease Ontology. When an ancestor term and its child terms have the same orthogroups, we remain only the ancestor term. If several terms with the same orthogroups share a common ancestor on the list, we also removed those terms to avoid using excessively detailed disease terms.

### Mapping of Gene Ontology terms considering the Gold Standard training set

We retrieved GO terms assigned for each human gene from the UniProt database^99^. Based on these human genes, we mapped vertebrate-level orthogroups for each GO term. Similar to Disease Ontology terms, we inherited orthogroups of ancestor terms to their child terms. Among the three categories of GO terms, we used GO cellular component (GOCC) and biological process (GOBP) terms.

Especially for GOCC terms, we separated terms based on their top ancestors: protein-containing complex, virion component, and cellular anatomical entity. Our data had no term whose top ancestor was the virion component and had assigned orthogroups. Next, we divided the protein-containing complex and cellular anatomical entity terms into terms that highly overlapped with the Gold Standard training sets and less overlapped ones. To determine the overlap between GOCC terms and the training sets, we calculated the Jaccard Index between one GOCC term and every training set and used the maximum value for each GOCC term. 0.5 was used as the threshold between high and low overlaps.

### Validating interactions by immunoprecipitation / mass spectrometry

All IP-MS experiments were conducted using the Pierce Co-Immunoprecipitation Kit (Thermo Scientific, #26149), according to the manufacturer’s instructions. Additional information on antibody identification is provided in **Table S1**.

Following elution of IP-MS, samples were resuspended in 50 μl of 2% SDS/50mM NH_4_HCO_3_ solution. Samples were heated at 95 °C for 5 minutes on a heat block to dissolve any visible tissues, immediately followed by overnight precipitation with 300μl precooled acetone in 4 °C. On the next day samples were centrifuged at 13,000 rpm (Eppendorf, 5430R) to pellet proteins, and the supernatant was discarded. The pellets were washed twice with 400μl precooled acetone, followed by the same centrifugation and supernatant aspiration steps. To resuspend protein pellets back to solution, to each sample 50μl 1% sodium deoxycholate (SDC)/50mM NH_4_HCO_3_ were added and sonicated twice in water bath (10 min. each). Samples were reduced with 5mM TCEP (Thermo Scientific, REF #77720) at 56 °C for 45 minutes, alkylated with 25mM iodoacetamide (Sigma, I1149-5G) in the dark at room temperature for 45 minutes, reactions were quenched with 12mM DTT subsequently. Proteins were digested overnight at 37℃ using 2μg trypsin (Thermo Scientific, REF #90058) per sample. After protein digestion, samples were acidified with 1% formic acid, precipitated SDC was removed by centrifugation of samples at 16,000 g for 10 minutes. Flowthrough was brought to the Amicon 10kDa filter (Millipore Sigma, UFC501024), and desalted using MilliporeSigma™ ZipTip™ Pipet Tips (Fischer Scientific, Catalog No. ZTC18S096), dried in speedvac and stored at −20 °C until further use for mass spectrometry.

Specific protein interactors (“preys”) were identified by proteomics from biological replicate experiments with at least 3 replicate mass spectrometry analyses (technical replicates, conducted on the Orbitrap Lumos instrument as detailed above) each, identifying proteins using ProteomeDiscoverer/Percolator and maintaining FDR<1% at the PSM, peptide, and protein level. Experiments were accompanied with paired negative control IP experiments conducted using antibodies targeting either GFP or the FLAG epitope tag (neither of which was present in the samples) in order to control for non-specific enrichment of proteins during the immunoprecipitation protocol. Significant interaction partners were identified using the software tool Degust^100^ and modeling differential enrichment between experimental and control IP-MS experiments using the edgeR quasi-likelihood method^101^ in concert with the remove unwanted variation (RUVr) method^102^ for the normalization of peptide spectral counts between samples, selecting differentially enriched proteins with a false discovery rate of <1%. IP-MS experiments in which the known target of the antibody was not significantly enriched were discarded prior to subsequent analyses.

### Summary of RELCH mouse data, sourced from KOMP/IMPC

With permission from the Knockout Mouse Program (KOMP), we sourced the RELCH mouse knockout data, which can be accessed through the website of the International Mouse Phenotyping Consortium (IMPC). Phenotype data collection procedures can be accessed in detail at https://www.mousephenotype.org/impress/index

### Zebrafish husbandry, genetics, and imaging

#### Zebrafish Maintenance and Care

All zebrafish experiments were performed according to University of Texas at Austin IACUC standards. Wild-type AB strains were used unless otherwise stated. Embryos were raised at 28.5 ℃ in fish water (0.15% Instant Ocean in reverse osmosis water) and then transferred to standard system water at 5 dpf. Both male and female zebrafish were used for all adult assessments. As zebrafish do not exhibit sexual dimorphism until the early adult stages of development, sex is not a factor in any of the embryonic assessments described in this manuscript.

#### Generation of the arhgef1bdf allele

We generated a deficiency in the *arhgef1b* locus by co-injecting one-cell stage embryos with CRISPR sgRNAs targeting the 5’– and 3’-ends of the gene. Specifically, CHOPCHOP v3^103^ was used to design sgRNAs targeting exon 2 near the start site (5’-TGGTAGGACCTGCGACAGCA-3) and exon 30 near the stop site (5’-GCTAAATACACACTTCAGGG-3’) of the *arhgef1b* locus. The sgRNAs were acquired from Synthego with modifications to contain 2’-O’Methyl at the first three and last three bases and 2’-phosphorothioate bonds at the first three and last two bases. One-cell stage embryos were injected with 1 nL of 5 µM equimolar sgRNA-Cas9 protein in 0.1 M KCl. For F0 notochord imaging, we injected into the transgenic vacuolated cell membrane marker line, *Tg(col8a1a::GFP-CAAX)*^104^. Once the F0 fish reached sexual maturity, sperm was collected from F0 males^105^ to screen for deficiency founders (see below). F0 founders were outcrossed to the *Tg(col8a1a:GFP-CAAX)* line, and F1 carriers were identified using the same genotyping method on gDNA from finclip samples. F1 heterozygous deficiency carriers were in-crossed for all imaging experiments.

#### DNA extraction from sperm, finclip, and whole embryo samples

DNA was extracted from sperm, finclip, and whole embryos using the HotShot method^106^. Briefly, samples were lysed in 50 mM NaOH and heated to 95 ℃ for either 40 (sperm) or 20 minutes (finclip and whole embryos) before neutralization with a 1:4 volume of 1 M Tris-HCl (pH 8.0). Samples were either immediately used for PCR amplification or were stored at –20 ℃ until further processing.

#### Single PCR genotyping method for identification of arhgef1bdf allele

Carriers of the *arhgef1b^df^* allele were identified by PCR using a forward primer upstream of the 5’-sgRNA cut site (5’-CTTGGCTTCTGGCAGTATTTTT-3’) and a reverse primer downstream of the 3’-sgRNA cut site (5’-CGACTCCAGCACATACAGTGAT-3’). Importantly, this primer pair does not produce an amplicon in the wild-type condition; therefore the presence of an amplicon indicates individuals carrying at least one copy of the *arhgef1b^df^*allele. All PCR reactions were performed using GoTaq Green Master Mix (Promega #M7123) and the following thermocycler settings: an initial denaturation at 95 ℃ for 3 minutes, followed by 34 cycles of denaturation at 95 ℃ for 30 seconds, annealing at 55 ℃ for 30 seconds, and extension at 72 ℃ for 30 seconds, followed by a final extension at 72 ℃ for 5 minutes. Amplicons were run on 2% agarose gels for 20 minutes.

#### Single PCR genotyping method for identification of wild-type arhgef1b allele

To distinguish between PCR failure and the absence of the deficiency allele, we designed a genotyping method to identify the wild-type *arhgef1b* allele. For this we used primers surrounding the 5’-sgRNA cut site (the same upstream 5’-TGGTAGGACCTGCGACAGCA-3’ primer and a new downstream 5’-ACCAAATCAGATACAGGCCACT-3’ primer). Importantly, this primer pair does not produce an amplicon in the homozygous *arhgef1b^df^* condition; therefore, the presence of an amplicon indicates individuals carrying at least one copy of the wild-type *arhgef1b* allele.

#### Duplex PCR genotyping method for distinguishing homozygous and heterozygous arhgef1bdf carriers

To distinguish wild type, heterozygous *arhgef1b^df/+^*, and homozygous *arhgef1b^df/df^* mutants in a single PCR reaction, we developed a duplex genotyping method. Since both deficiency and wild-type PCR reactions used the same forward primer (5’-TGGTAGGACCTGCGACAGCA-3’), we combined this with half-molar concentrations of each of the reverse primers (5’-CGACTCCAGCACATACAGTGAT-3’ and 5’-ACCAAATCAGATACAGGCCACT-3’). The PCR products were run on 4% agarose gels for at least one hour. Wild-type *arhgef1b* amplicons ran around 250 bp, while *arhgef1b^df^* amplicons ran slightly higher on the gel, and the presence of both bands indicated a heterozygous *arhgef1b^df/+^* individual.

#### Confocal microscopy and analysis

F0 *arhgef1b* or stable F2s (from F1 *arhgef1b^df/+^; Tg(col8a1a::GFP-CAAX)* in-crosses) were raised until 5 dpf, at which point they were anesthetized in 0.0016% Tricaine and mounted laterally in 1% low-melt agarose for imaging. Whole notochord stacks were acquired on the Nikon CSU-W1 Yokogawa Confocal Microscope and cell segmentation was performed in IMARIS (v10.2.0) as described in Voigt *et al*., 2025^91^. Briefly, all images were taken at 20X magnification with a 1.2 µm step size, and cell segmentation was performed using the IMARIS Cells and Surfaces modules with the following settings: Source channel 488 nm, cell smallest diameter of 20 µm, membrane detail of 2.0 µm, local contrast, and manual thresholding. All volume and distance datasets were exported and loaded into R for statistical analysis. Representative images were taken in IMARIS and colored by volume measurements.

## ACKNOWLEDGEMENTS

The authors gratefully acknowledge the contributions of the Knockout Mouse Program (KOMP2) at the Baylor College of Medicine and the International Mouse Phenotyping Consortium (IMPC) for the *Relch^em1(IMPC)J^* mice data, Wayne McFee at NOAA for providing dolphin brain samples, Juju Dessert for animal and brain cell figure illustrations, John Wallingford for manuscript feedback, Kevin Drew and Anna Battenhouse for extensive assistance with computational analyses, Maria Person, Michelle Gadoush, and Peter Faull at the UT Proteomics Facility for aiding in data collection, and the Texas Advanced Computing Center at The University of Texas at Austin for providing high-performance computing resources. Research was funded by grants from the National Institute of General Medical Sciences (R35GM122480 to E.M.M.), the National Institute of Arthritis & Musculoskeletal & Skin Diseases (R01AR071009 to R.S.G.), the National Institute of Child Health and Human Development (F31HD114419 to B.V.), the United States Army Research Office (W911NF-12-1-0390 to E.M.M.), and the Welch Foundation (F-1515 to E.M.M.). Identification of certain commercial equipment, instruments, software, or materials does not imply recommendation or endorsement by the National Institute of Standards and Technology, nor does it imply that the products identified are necessarily the best available for the purpose.

## DATA AVAILABILITY

Detailed Supplemental Methods are provided, including 7 supplemental figures and 5 supplemental tables. All raw and interpreted mass spectrometry data were deposited to the ProteomeXchange *via* the MassIVE partner repository with the identifiers provided in **Table S1**. A key resource table containing all resources used in this study, including github links for code, is deposited at Zenodo (doi:10.5281/zenodo.14807396).

## SUPPLEMENTAL FIGURES

**Figure S1.**
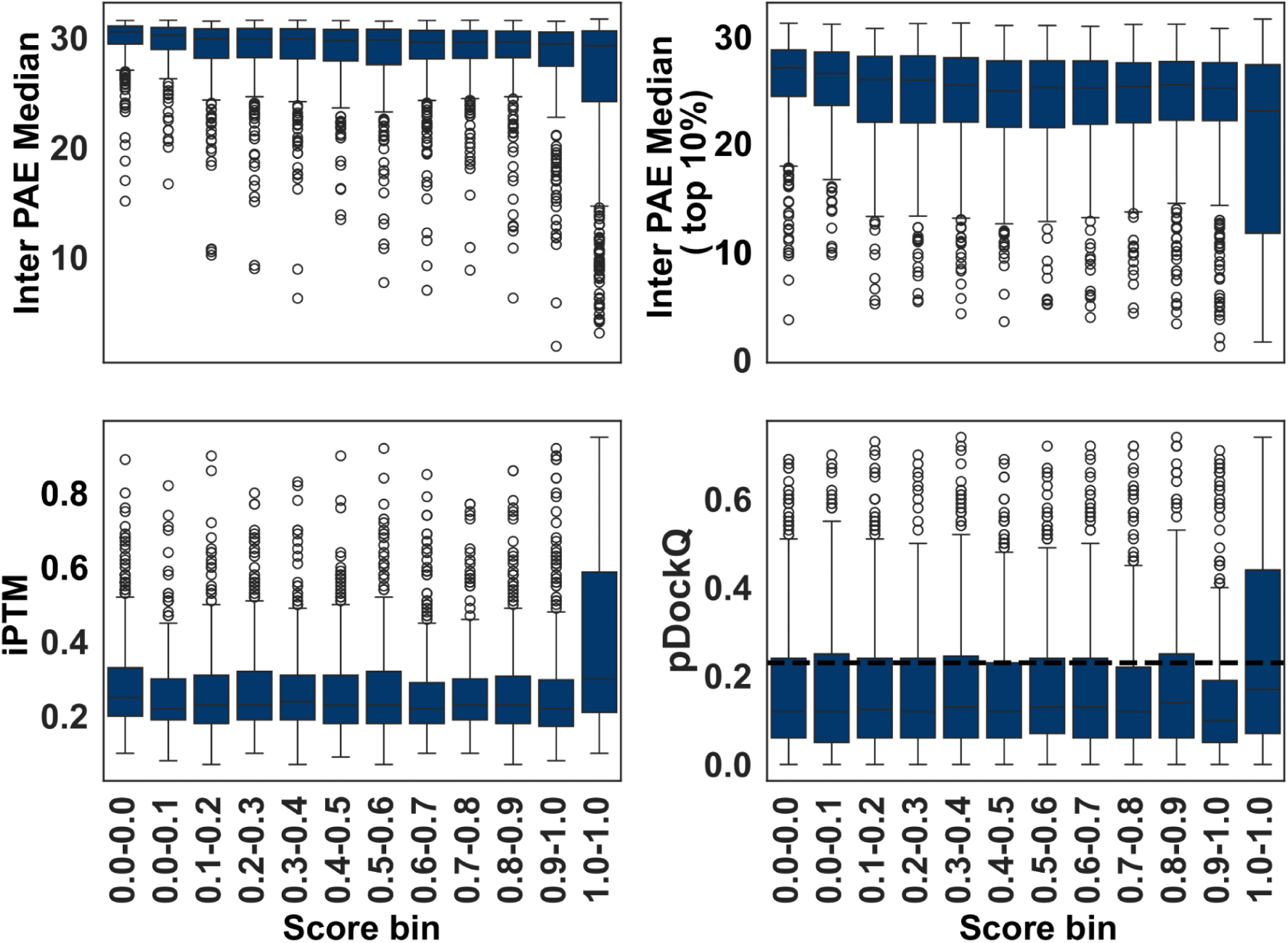
Alphafold-multimer score distributions for sampled protein pairs as a function of CF-MS interaction score.

**Figure S2.**
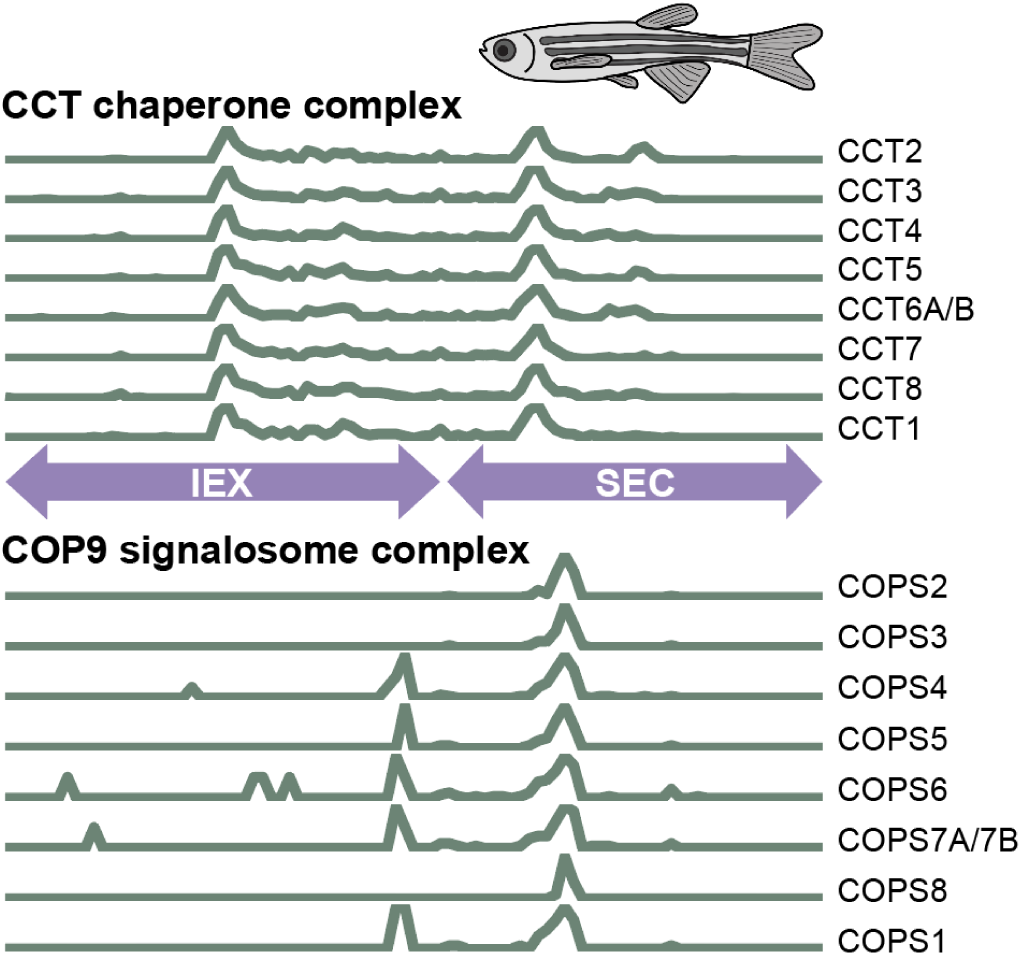
Co-elution profiles show intact subunits of the CCT chaperone and COP9 signalosome complexes from CF-MS experiments in zebrafish brains. Elution profiles were derived from ion exchange chromatography (IEX) followed by size exclusion chromatography (SEC). Each line represents the abundance of an individual complex subunit of the CCT chaperone and COP9 signalosome complexes across chromatographic fractions. The consistent co-elution of all annotated subunits within each complex indicates intact assembly and validates the resolution and reliability of the CF-MS workflow in adult zebrafish brain lysates.

**Figure S3.**
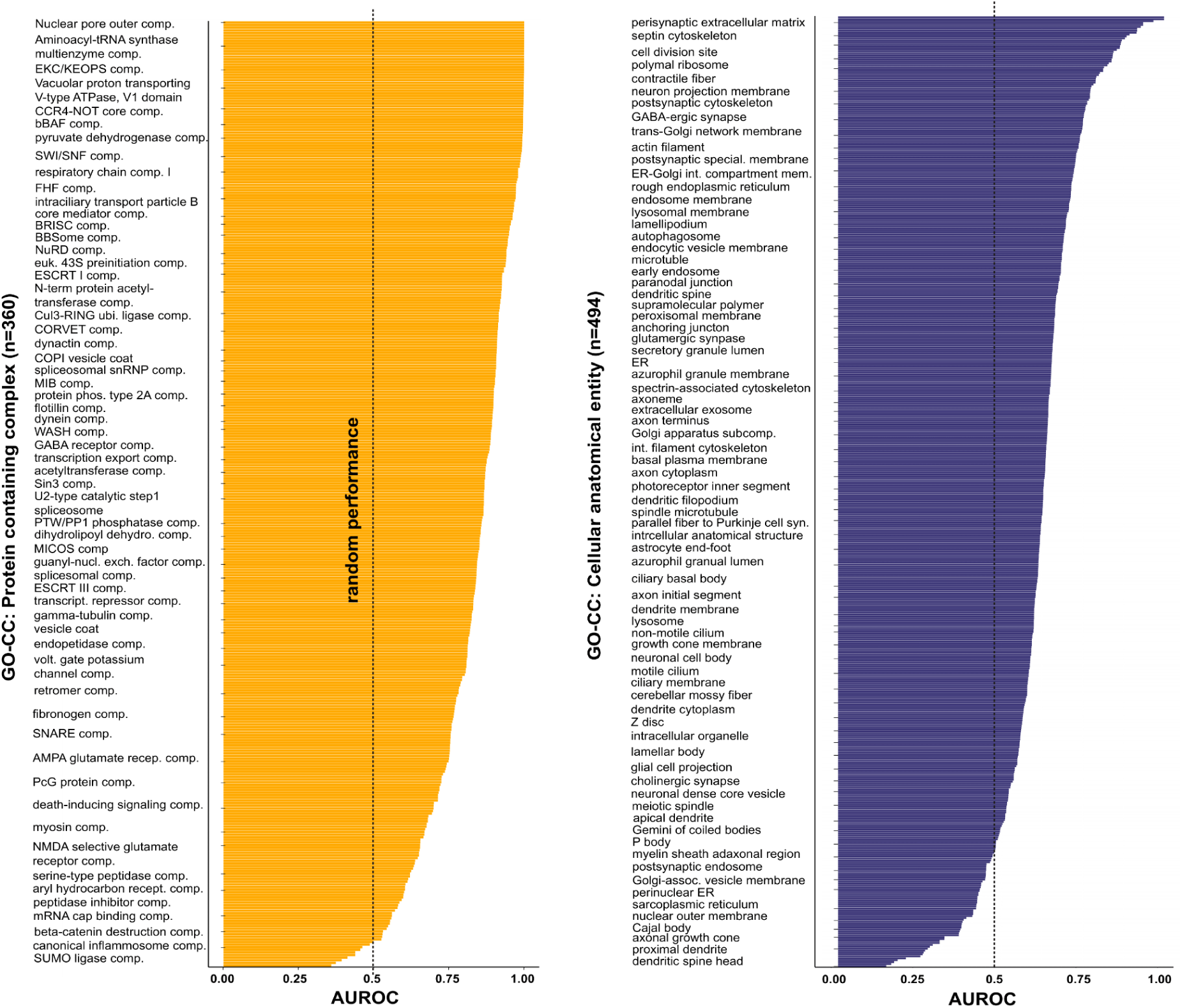
Area under the receiver operating characteristic curve (AUROC) values for correctly associating proteins in VerteBrain that are known to belong to the same Gene Ontology (GO) Protein containing complex or Cellular anatomical entity. Leave-one-out network propagation analysis (see **Methods**) was used to evaluate how well protein-protein interaction scores from the VerteBrain dataset recapitulate known protein co-memberships in Gene Ontology (GO) categories. Left: AUROC values for proteins annotated as part of the same GO cellular component term categorized as a “Protein-containing complex” (n = 360 terms). Right: AUROC values for GO terms categorized as “Cellular anatomical entity” (n = 494 terms). High AUROC values indicate strong agreement between VerteBrain interactions and established GO co-complex or co-localization annotations. Dotted vertical lines indicate the baseline of random performance (AUROC = 0.5)

**Figure S4.**
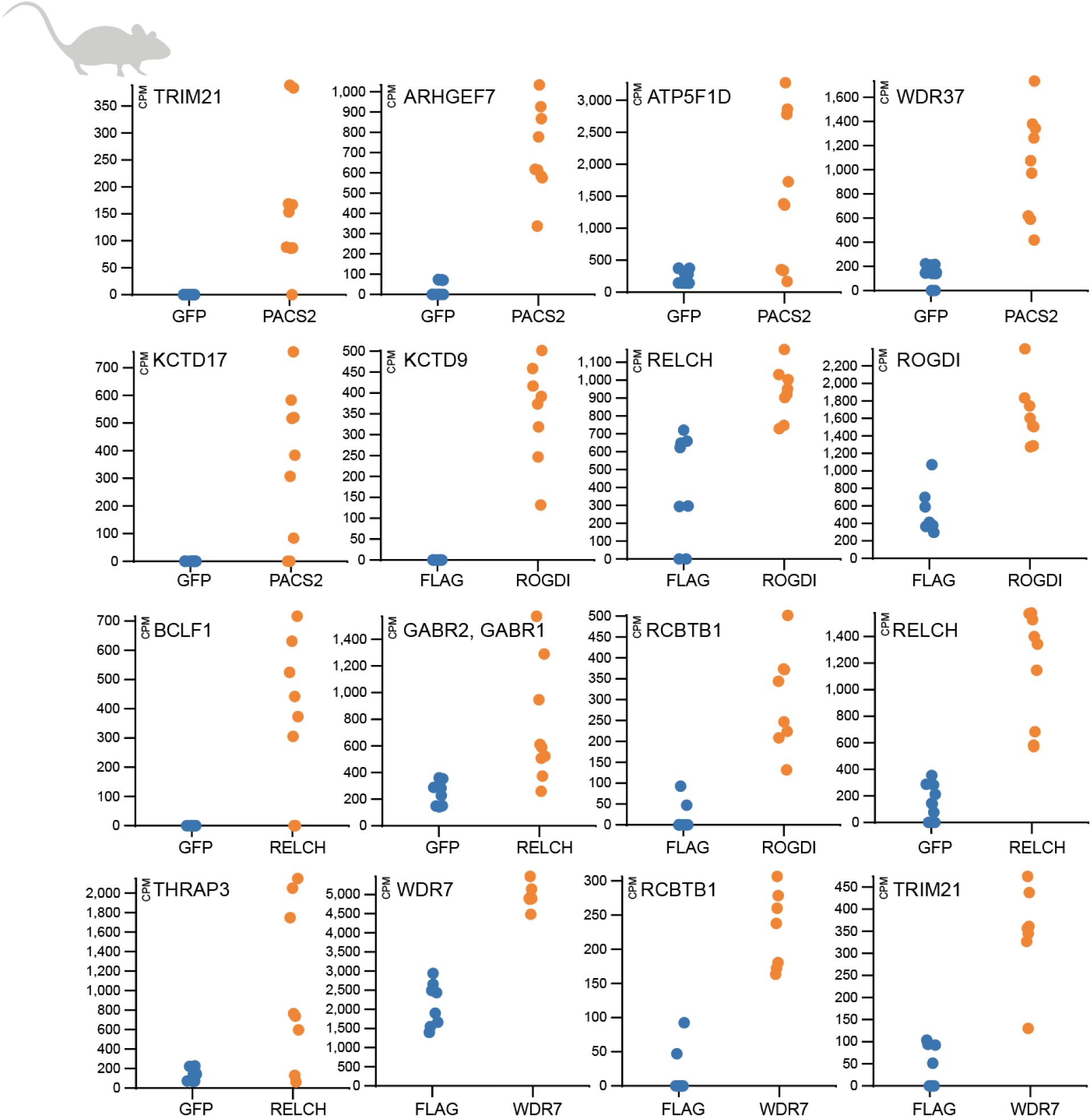
Enrichment plots for selected significant (FDR < 1%) interactors identified from IP-MS experiments in mouse whole brain lysates. Each plot reports mass spectrometry-measured abundances (colored dots) of the prey protein labeled in the top left of the plot following immunoprecipitation from whole mouse brain with antibodies targeting the bait protein labeled on the x-axis, accompanied by data from matched IP experiments using antibodies targeting either GFP or FLAG as negative controls. Abundances are reported in normalized counts per million (cpm) as measured using Degust (see **Methods**) across 3-4 replicate mass spectrometry analyses of each biological replicate pulldown.

**Figure S5.**
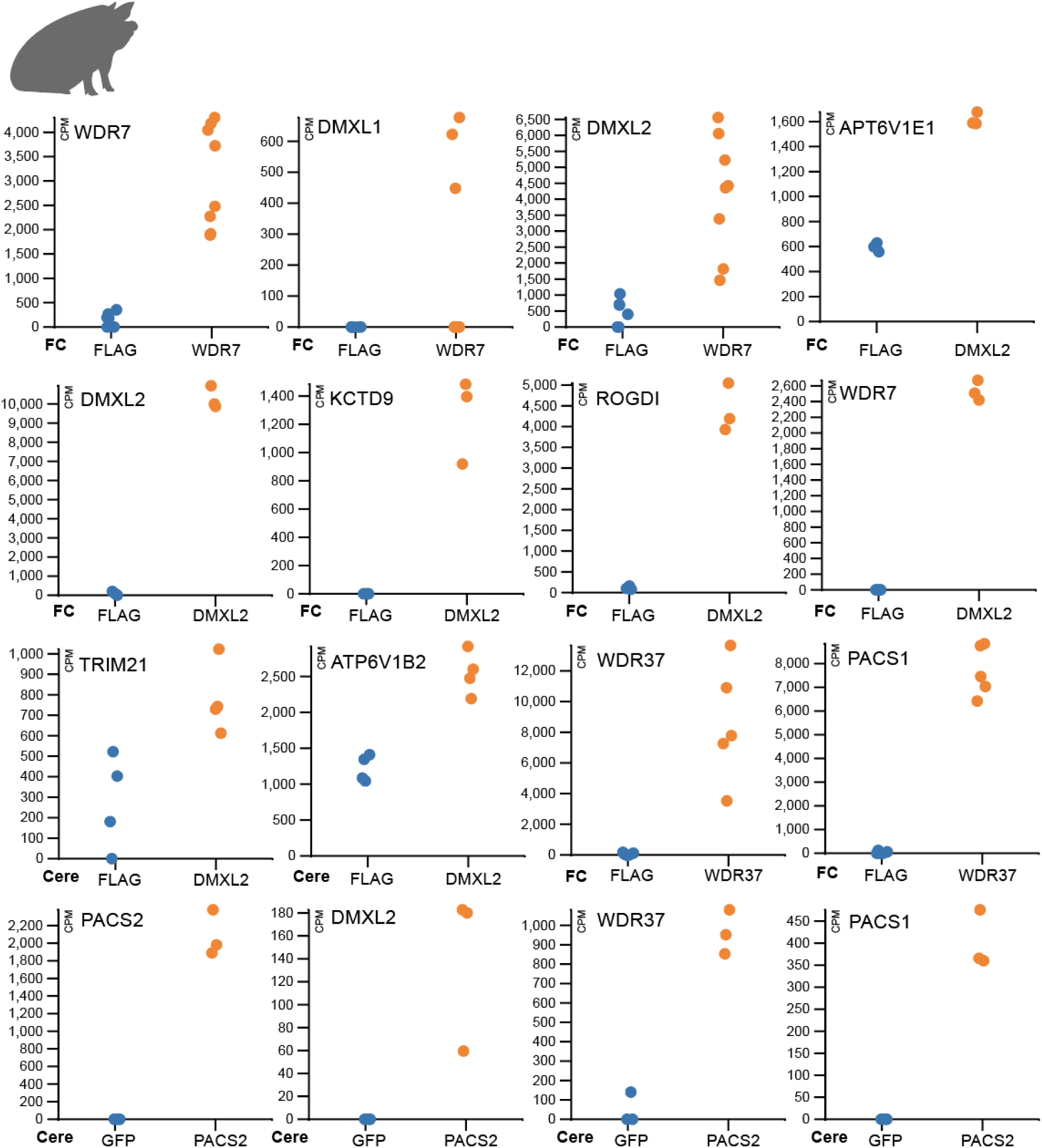
Enrichment plots for selected significant (FDR < 1%) interactors identified from IP-MS experiments in pig brain lysates. Each plot reports measured mass spectrometry-measured abundances (colored dots) of the prey protein labeled in the top left of the plot following immunoprecipitation from pig brain (either frontal cortex (FC) or cerebellum (cere), as noted) with antibodies targeting the bait protein labeled on the x-axis, accompanied by data from matched IP experiments using antibodies targeting either GFP or FLAG as negative controls. Abundances are reported in normalized counts per million (cpm) as measured using Degust (see **Methods**) across 3-4 replicate mass spectrometry analyses of each biological replicate pulldown.

**Figure S6.**
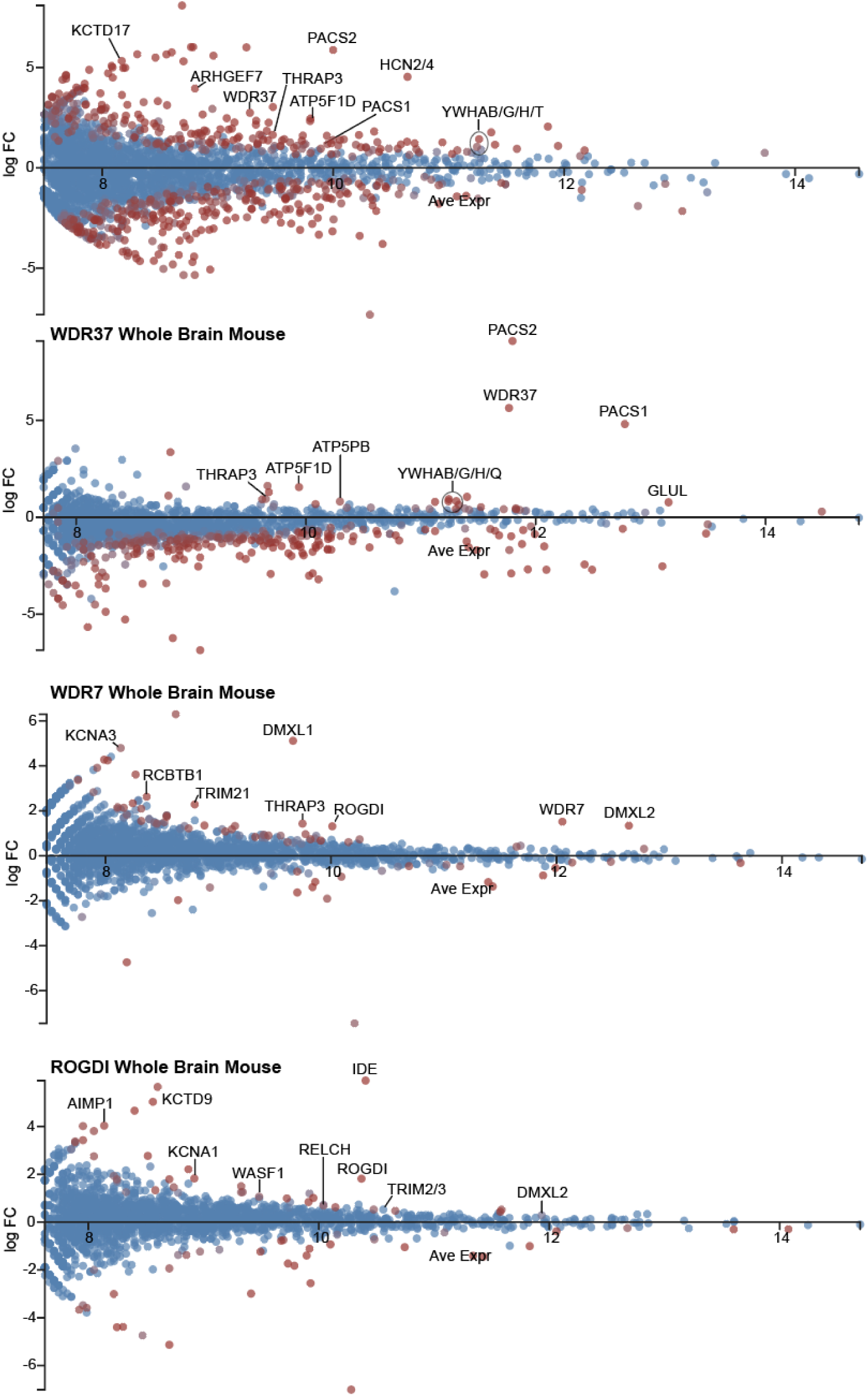
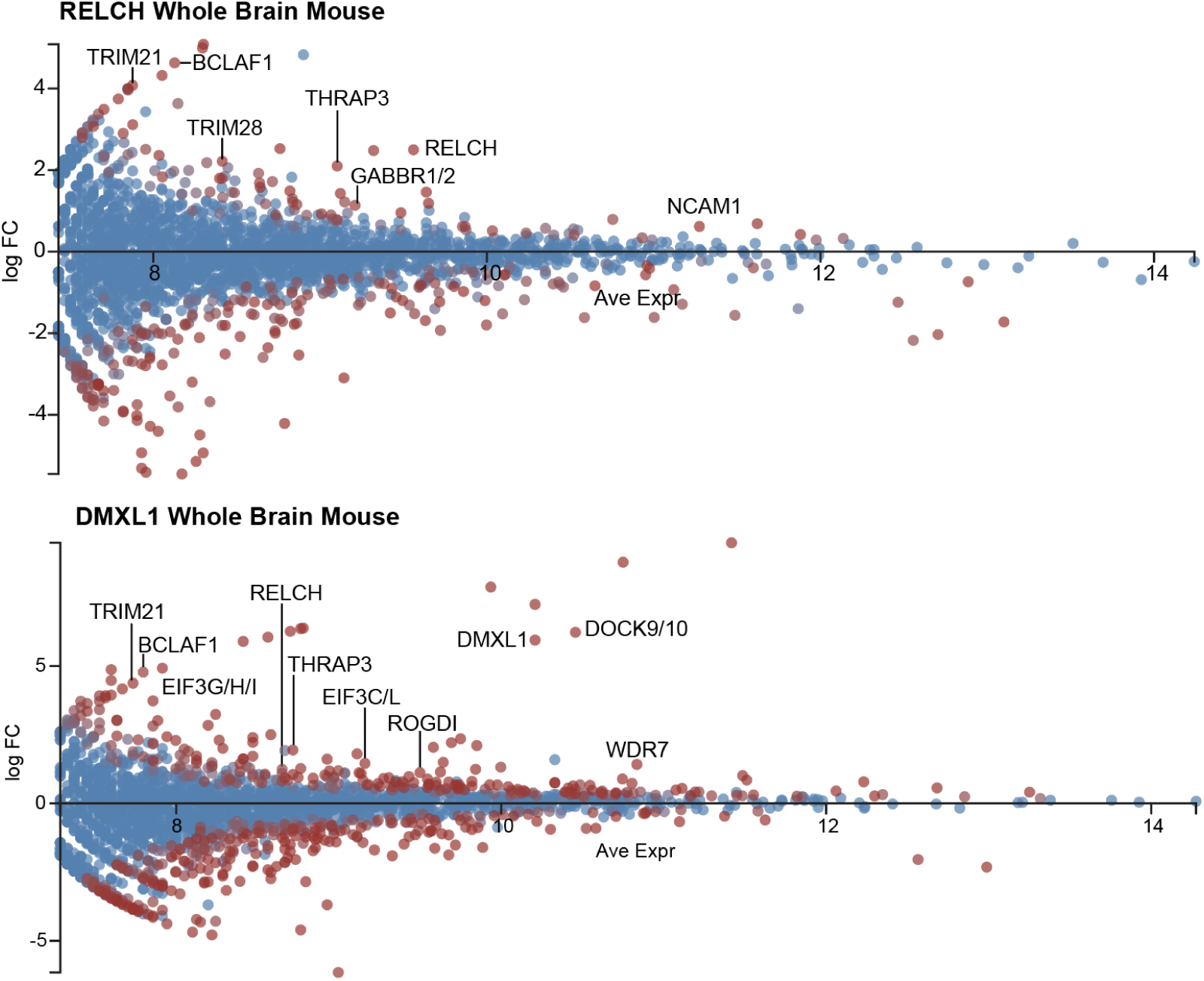
MA plots for selected IP-MS experiments in mouse whole brain lysates. Each plot reports the log mean abundances (x-axis) versus log ratios (y-axis) of all proteins (colored dots) detected by mass spectrometry at confidence (FDR < 1%) following immunoprecipitation from whole mouse brain with antibodies targeting the bait protein labeled at the top of the plot or by matched IP experiments using antibodies targeting either GFP or FLAG as negative controls. Proteins with positive log ratio values are enriched in the experimental samples; proteins with negative log ratio are enriched in the negative control IPs. Proteins are colored on a blue (non-significant) to red (significantly differentially enriched, FDR < 1%) color scale. Abundances are reported in normalized counts per million (cpm) measured using Degust (see **Methods**) and are matched to the plots in **Figure S4**.

**Figure S7.**
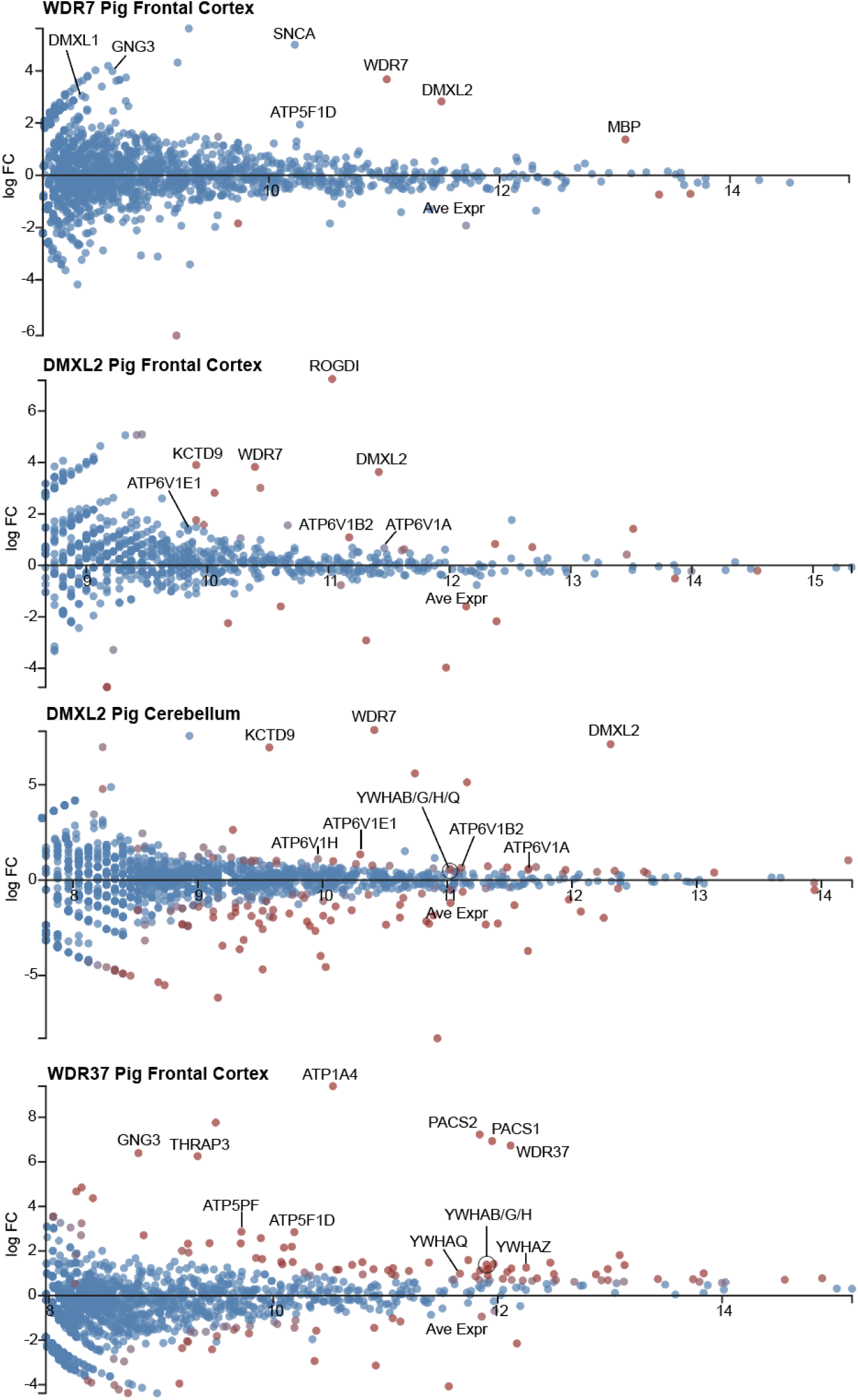
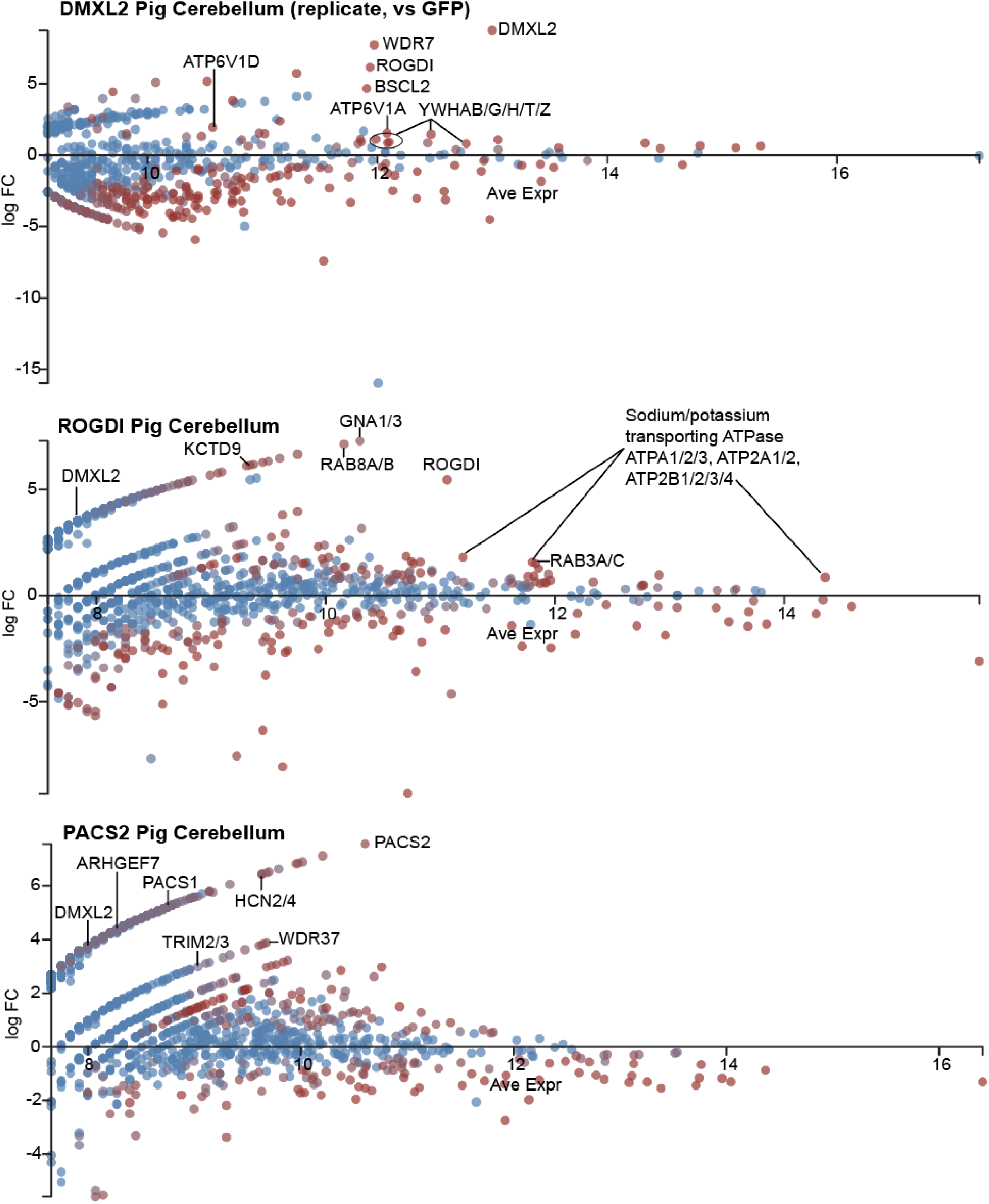
MA plots for selected IP-MS experiments in pig brain lysates. Each plot reports the log mean abundances (x-axis) versus log ratios (y-axis) of all proteins (colored dots) detected by mass spectrometry at confidence (FDR < 1%) following immunoprecipitation from pig brain with antibodies targeting the bait protein (labeled at the top of the plot along with the tissue analyzed) or by matched IP experiments using antibodies targeting either GFP or FLAG as negative controls. Proteins with positive log ratio values are enriched in the experimental samples; proteins with negative log ratio are enriched in the negative control IPs. Proteins are colored on a blue (non-significant) to red (significantly differentially enriched, FDR < 1%) color scale. Abundances are reported in normalized counts per million (cpm) measured using Degust (see **Methods**) and are matched to the plots in **Figure S5**.

## SUPPLEMENTAL TABLES

**Table S1:** Overview summary of biological samples, reagents, software, and algorithms used in this construct the vertebrate brain interactome.

**Table S2:** Quantification in peptide-spectral matches of 9,259 verNOG orthogroups across 2,197 biochemical fractions from 35 CF-MS experiments using post-mortem brain tissue samples in five species, related to Figure 1.

**Table S3:** Pairwise interactions in VerteBrain above 8% FDR threshold based on the ExtraTreeClassifier model.

**Table S4:** 6,108 verNOG orthogroups organized hierarchically into varying granularities of protein assemblies, denoted by the column headers, e.g. as for the column headed “cut_3926”, which provides 3,926 protein assemblies, identified by numerical cluster identifiers.

**Table S5:** Summary of significant interactors measured by IP-MS of Rabconnectin-3 and WDR37/PACS subunits relative to control IP-MS experiments.

